# Long-term reliable neural decoding based on flexible implantable microelectronics and machine learning for seizure prediction application

**DOI:** 10.1101/2023.03.08.531452

**Authors:** Zicong He, Jiajun Zheng, Junwei Duan, Zhe Jin, Zixuan Huang, Shuaishuai Wu, Qian He, Kwok-Fai So, Shuixing Zhang, Zhiyuan Xiong

## Abstract

Neural decoding is useful for understanding brain functions and developing neural interface applications. However, neural interfaces based on rigid electronics often suffer from recording instability due to the foreign body responses caused by their mechanical mismatch with soft tissues, limiting the longitudinal accuracy of neural decoding methods. Herein, it is reported that flexible electronics can be integrated with machine learning algorithms to achieve long-term reliable neural decoding. Wet-spun conductive polymer microfibers showed mechanical robustness and flexibility, low impedance, and chronic biocompatibility, enabling intracerebral neural recordings in epileptic mice at a high signal-to-noise ratio eight weeks after implantation. When the signals recorded by the flexible electrodes were used in machine learning analyses with diverse complex algorithms, they consistently showed higher prediction accuracy for epileptic seizures than stiff metal electrode signals, particularly in the case of using long-term recordings for testing or small-sample datasets for training. A real-time warning system based on the flexible neural electrodes was built that predicted seizures eight minutes in advance with a low false alarm rate. Our work bridges flexible electronics and artificial intelligence for neural decoding applications such as long-term treatment of chronic neurological disorders.

## Introduction

The field of neural decoding explores the use of signal processing methods to decipher the mechanisms of neural activity to understand brain functions and develop various neural interface applications, such as restoring body functions (*e.g.*, prosthesis control, sensation restoration, language output and cognitive enhancement)^1, 2^ and treating neurological diseases (*e.g.*, seizures, Parkinson’s, Alzheimer’s, and mood disorders)^1, 3^. In this context, the decoding algorithms and the neural interfacing techniques that generate neural signals as the input to the decoders are highly intertwined to collectively impact the decoding performance^4–7^. Generally, a high-resolution neural recording interface enables the decoder to automatically select informative aspects of the recordings to build relationship with the physiological quantities of interest^8–10^, and can be used to implement an informed decoder that utilizes the understanding of neural coding principles^11^. However, existing neural interface techniques, such as electroencephalogram, electrocorticogram and microelectrode arrays, often balance the recording resolution with the long-term stability due to foreign body responses such as chronic inflammation and glial scarring^12, 13^. Recording instability leads to degradation or fluctuations in the decoding performance over time^5, 14^ and necessitates time-consuming recalibration for implantable brain-computer interface (BCI) devices^15, 16^. This instability can be partially solved by adopting adaptive decoders; however, this increases the computational costs^16, 17^, and a universal strategy to realize long-term reliable neural decoding is still lacking.

Previous research has shown that one key reason for the foreign body responses is the significant mechanical mismatch between the neural recording electrodes based on stiff metal materials and the soft neural tissues^18, 19^. This mismatch generates strong mechanical stress stimuli on the interfaced neural cells when micromotion of the electrodes occurs, resulting in shear-induced inflammation or encapsulation of the electrodes^20, 21^. With the rapid development of materials and nanotechnology, flexible neural electrodes have been designed to minimize the electrode/tissue mechanical mismatch^22–25^. In the past decade, flexible neural interfaces have been fabricated with new electroactive materials (*e.g.*, conductive polymers^18, 26, 27^, carbon nanotubes^28^, and graphene^29–31^) and structural engineering strategies (*e.g.*, gelation^18, 26, 32, 33^ and miniaturization^34–36^). Compared with conventional neural interfaces, flexible neural interfaces show unique capabilities to access diverse neural structures (*e.g.*, brain^37^, spinal cord^38^, and peripheral nerves^39^) and monitor various neural activities (*e.g.*, single-unit spiking^27, 35^ and local field potentials^13^) with tuneable recording resolution and chronic stability over months to even years^18, 26, 38, 40–42^. Thus, flexible electronics appear to be an ideal choice to pursue accurate yet stable neural decoding. However, current works in flexible neural electrodes have been mostly devoted to the front-end signal recordings but less attention has been given to back-end data processing. To explore the potential of flexible electronics in neural decoding, the interplay between this new interface technique and the decoding algorithms in scenarios with varied volume/quality of the recorded neural signals as well as different algorithm complexity, and particularly how their interplay evolves over time to influence the decoding stability need to be examined.

In this work, we developed conductive polymer microfibers as flexible neural microelectrodes and explored their integration with machine learning algorithms for use in neural decoding. The polymeric microelectrodes combine superior mechanical flexibility and electrochemical properties, thereby enabling biocompatible neural interfaces for precise and robust decoding of neural activities related to disease states. This work provides new opportunities for the use of flexible electronic techniques for neural decoding in clinically relevant applications.

## Results

### Preparation and characterization of flexible conductive polymer fibers

Poly(3,4-ethylenedioxythiophene):poly(styrene sulfonate) (PEDOT:PSS), a conventional and commercially available conductive polymer, was chosen to fabricate the flexible neural electrodes owing to its good solution processability and wide use as bioelectronic materials^43–47^. PEDOT:PSS dispersions were fabricated into fibers through the widely used wet spinning technique. First, the PEDOT:PSS dispersions were injected into a sulfuric acid-based coagulation bath (Fig. 1a), which removed the PSS and water to induce crystallization of the PEDOT chains^47–49^, resulting in the formation of highly elastic hydrogel fibers (Fig. S1). Next, the elastic hydrogel fibers were washed and dried under drawing stress (Fig. 1a), generating free-standing PEDOT microfibers (Fig. 1b) with ribbon-shaped cross sections and varied lateral dimensions (Figs. 1c and S2). The wide-angle X-ray scattering measurements showed that the drawing stress promoted the PEDOT laminar crystals to be oriented along the fiber axis (Figs. 1d and S3), leading to both high tensile strength (3216.1 ± 284.9 S cm^-1^) and electrical conductivity (428.4 ± 81.7 MPa) for the resultant fibers (Fig. S4). These values are 3−4 times larger than the values (1059.1 ± 337.3 S cm^−1^ and 111.7 ± 17.5 MPa) of the naturally dried fibers (Fig. S4) and are very competitive among the previously reported PEDOT fibers that were wet-spun under different conditions (Fig. 1e)^48–59^. The high strength of the conductive PEDOT fibers facilitated the implantation operation.

**Fig. 1.**
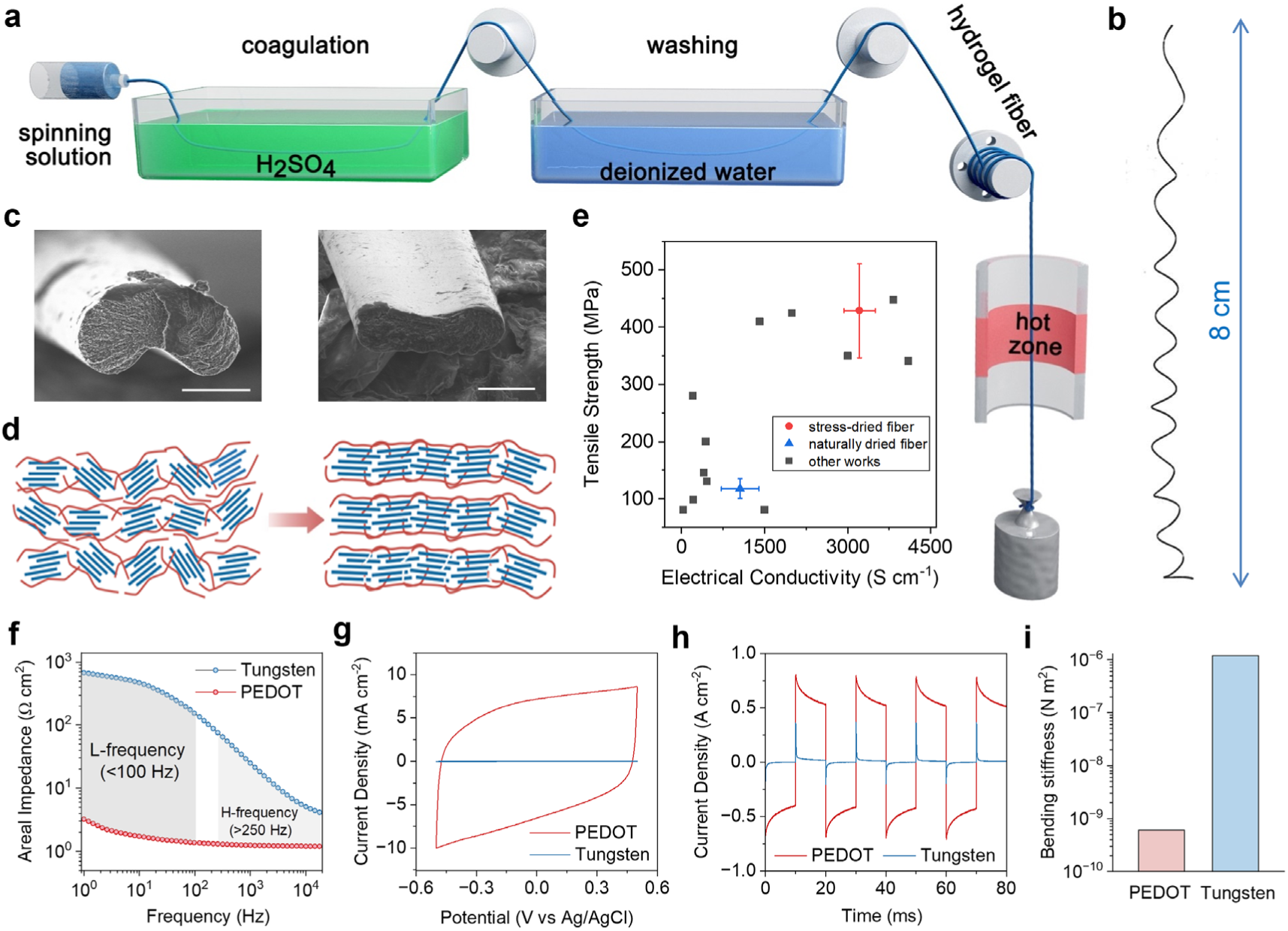
Fabrication and characterization of the PEDOT fibers. (**a**) Schematic illustration of the fabrication process of the PEDOT fibers through wet spinning. (**b**) A photograph of a freestanding PEDOT fiber coil. (**c**) Scanning electron microscope images of the cross section of the PEDOT fibers. Scale bar: 30 μm. (**d**) Schematic diagram of the enhanced orientation of the PEDOT microcrystals during drying under stress. The blue and red lines represent the PEDOT and PSS chains, respectively. (**e**) Comparison of the tensile strength and electrical conductivity of naturally dried, stress-dried, and previously reported PEDOT fibers. (**f**-**g**) Electrochemical impedance spectra and cyclic voltammetry curves of the PEDOT fibers and tungsten wires. (**h**) Current measurement after bipolar pulse voltage stimulation of the PEDOT fibers and tungsten wires. (**i**) Comparison of the bending stiffnesses of the PEDOT fibers and tungsten wires. The PEDOT fibers used in Fig. 1 were prepared using 13 mg mL^−1^ PEDOT:PSS dispersions as spinning dopes and 21G needles.

We evaluated the electrochemical performance of the PEDOT fibers as neural recording electrodes. Clinically adopted neural electrodes based on tungsten wires with similar lateral dimensions (80 μm diameter) to the PEDOT fibers (70−80 μm width) were used as the control. The electrochemical impedance of the PEDOT fiber electrodes in phosphate-buffered saline (PBS) was found to be 1−2 orders of magnitude lower than that of the tungsten wire electrodes in the high-frequency region of > 250 Hz (Fig. 1f). In particular, the difference was enlarged to nearly 3 orders of magnitude in the high-frequency region of <100 Hz, indicating that the PEDOT microelectrodes may be more sensitive at recording low-frequency neural activity, such as the local field potential studied in the following experiments. The small impedance of the PEDOT microelectrodes in the low-frequency region is attributed to the large interfacial capacitance^43^, as their charge storage capacity (Figs. 1g and S5) and charge injection capacity (Figs. 1h and S6) are significantly higher than those of the tungsten wire electrodes.

Impressively, the PEDOT fibers show a low bending stiffness of ∼6.0 × 10^−10^ N m^2^ owing to their ribbon-shaped cross section (see the detailed calculation in the Supporting Information). This bending stiffness is more than 3 orders of magnitude smaller than that (∼1.2 × 10^−6^ N m^2^) of the tungsten wires and is comparable to the stiffnesses of several state-of-the-art polymeric neural microelectrodes^60, 61^. The good mechanical flexibility of the PEDOT fibers was further confirmed by their excellent knittability (Fig. S7) and was visualized through a bulb lighting test, in which the bulb light could be maintained even when the PEDOT fiber-based connecting lines were bent at an angle of 180° (Video 1). Moreover, the electrochemical impedance and capacitance of the PEDOT fibers in PBS remained essentially unchanged after 10000 bending cycles (Fig. S8), 8 weeks of immersion (Fig. S9) and millions of pulse voltage stimulations (Fig. S10), indicating the superior stability of flexible PEDOT fiber electrodes for *in vivo* use.

### Chronic Biocompatibility and Stability

We next evaluated the performance of the PEDOT microfibers and tungsten wires, which represented flexile and stiff neural electrodes, in intracerebral neural recordings. A common drug-resistant epilepsy, temporal lobe epilepsy, was used as the experimental model, and the electrodes were insulated with Parylene-C and implanted into the mouse hippocampus to monitor normal and epileptic neural activity.

Magnetic resonance imaging (MRI) confirmed the successful implantation and 8-week stability of the PEDOT microelectrodes. As shown in Fig. 2b, the implanted microelectrodes did not exhibit any obvious displacement in the 8 weeks after surgery. Compared to the uninjured sides, there was no additional T_2_-weighted hyperintensity related to oedema or haemorrhage surrounding the microelectrodes. In addition, consistent alterations in the T_2_ signals were observed after further quantification of the bilateral cortex and hippocampus (Fig. 2c), suggesting the absence of significant additional damage at the implantation sites.

**Fig. 2.**
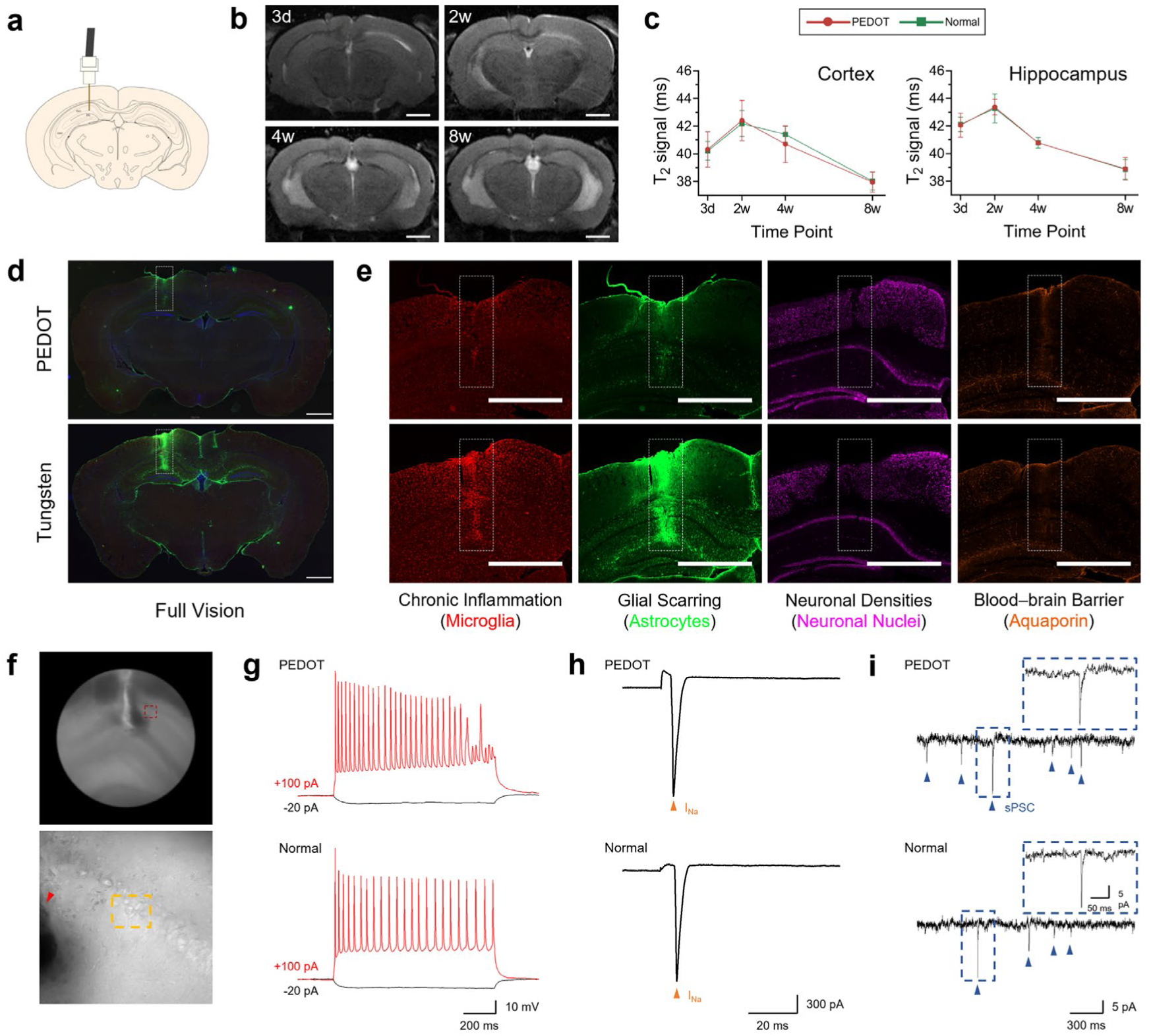
Chronic biocompatibility and stability. (**a**) A schematic diagram of the microelectrode implantation in the hippocampal CA1 region. The coronal section was modified from Paxinos and Franklin^62^. (**b**) Magnetic resonance (MR) images were acquired 3 days, 2 weeks, 4 weeks, and 8 weeks after PEDOT microelectrode implantation. Scale bar: 1000 μm. (**c**) T_2_ signal quantification in the bilateral cortex (*n* = 3 for each; student’s paired-samples t tests, *t*_3d_ = 0.116, *p* > 0.05; *t*_2w_ = 0.228, *p* > 0.05; *t*_4w_ = −1.006, *p* > 0.05; *t*_8w_ = −0.368, *p* > 0.05) and hippocampus (*n* = 3 for each; student’s paired-samples t tests, *t*_3d_ = −0.217, *p* > 0.05; *t*_2w_ = 0.240, *p* > 0.05; *t*_4w_ = −0.065, *p* > 0.05; *t*_8w_ = 0.153, *p* > 0.05). A similar signal decrease was observed in the bilateral cortices and hippocampi, which was the result of tissue atrophy caused by epilepsy. (**d**) The implantation injuries caused by the PEDOT (top) and tungsten (bottom) microelectrodes. Scale bar: 1000 μm. (**e**) Brain slices were analysed by immunohistochemistry staining with microglia for chronic inflammation (red), astrocytes for glial scarring (green), neuronal nuclei for neuronal densities (purple), and aquaporin for the blood-brain barrier (orange). Scale bar: 1000 μm. (**f**) The patch-clamp recordings were performed at the sites surrounding the implantation trajectory of the PEDOT microelectrodes. 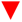 represents the implantation trajectory. (**g**-**i**) Electrophysiological measurements of the (**g**) action potential (AP), (**h**) voltage-gated sodium current (I_Na_), and (**i**) spontaneous postsynaptic current (sPSC) of neurons in the bilateral hippocampi.

Immunohistochemistry in the isolated brain tissues was performed 8 weeks after implantation to evaluate the foreign body response. As shown in the whole brain slices displayed in Fig. 2d, the implantation injury (marked by the box) in the PEDOT group was milder than that in the tungsten group. The details of the immunohistochemistry staining are shown in Fig. 2e. The accumulation of microglia and astrocytes was reduced at the implantation sites of the PEDOT microelectrodes compared with those of the tungsten microelectrodes, indicating an overall reduction in the extent of chronic inflammation and glial scarring. Correspondingly, the neuronal densities surrounding the PEDOT microelectrodes increased, indicating alleviation of neuronal degeneration. Moreover, the integral and organized blood-brain barrier labelled by aquaporin was clearly observed around the PEDOT microelectrodes, while it was disordered and incompact around the tungsten microelectrodes.

To study the influence of electrode implantation on the diversity and functions of the surrounding neurons, whole-cell patch-clamp recordings were performed in the bilateral hippocampal CA1 regions of living brain slices isolated at 8 weeks after implantation. As shown in Figs. 2g-i and S14, various electrophysiological properties, including the membrane capacitance, membrane resistance, action potential (AP) amplitude, voltage-gated sodium current (I_Na_) amplitude, and spontaneous postsynaptic current (sPSC) frequency and amplitude, showed no significant differences between neurons in the bilateral hippocampi. In addition, the presence of both excitatory and inhibitory neurons was confirmed at the implantation sides, which is consistent with the uninjured sides (Fig. S14).

### Intracerebral recordings of neural activity

In each group, we successfully observed 12−14 seizures in 6 mice during short-term recordings within 2 weeks postimplantation. The raw signals of whole cycles of epileptic seizures acquired by the PEDOT microelectrodes and their spectrograms converted by short-time Fourier transforms are shown in Figs. 3b, c. Epileptic activity occurred with gradually increasing discharges arising from the hyperexcitability and hypersynchrony of neurons. This activity was mainly associated with an increase in the spectral power in the low-frequency range between 0−30 Hz (Fig. 3c). After identifying the seizure event through the power amplitude, the epileptic seizure was divided into four states, including the interictal (the awake period between seizures), preictal (the period of approximately 10 minutes before seizure onset), ictal (the seizure period) and postictal (the period after seizure) states.

**Fig. 3.**
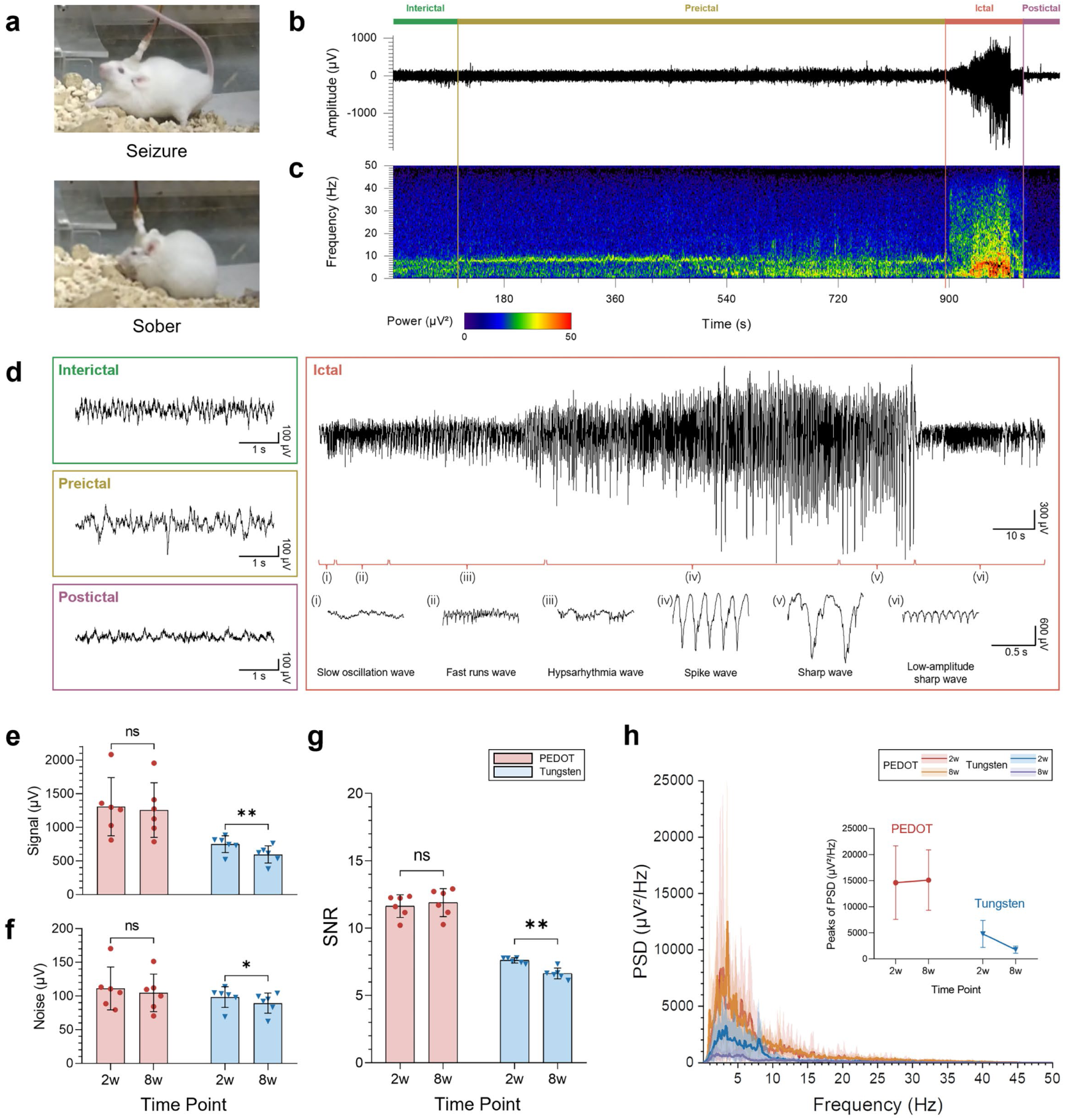
Intracerebral recordings of neural activities. (**a**) Neural recording in a temporal lobe epilepsy mouse with the PEDOT microelectrode. (**b**, **c**) The raw signals and their converted spectrograms for an epileptic seizure cycle acquired by the PEDOT fiber microelectrodes. (**d**) The signal details of four epileptic states. (**e**-**g**) The signal-to-noise ratio (SNR) comparison of epileptic signals between the short-term (0−2 weeks) and long-term (6−8 weeks) recordings for the PEDOT and tungsten groups, including the (**e**) signal amplitude (*n* = 6 for each; student’s paired-samples t tests, *t*_PEDOT_ = 1.733, *p* > 0.05; *t*_Tungsten_ = 5.778, *p* < 0.01), (**f**) noise level (*n* = 6 for each; student’s paired-samples t tests, *t*_PEDOT_ = 2.314, *p* > 0.05; *t*_Tungsten_ = 3.481, *p* < 0.05) and (**g**) SNR (*n* = 6 for each; student’s paired-samples t tests, *t*_PEDOT_ = −0.992, *p* > 0.05; *t*_Tungsten_ = 6.746, *p* < 0.01). * for *p* < 0.05, ** for *p* < 0.01, *** for *p* < 0.001 and *ns* for no significant difference. (**h**) Power spectrum density (PSD) analysis of epileptic signals between the short-term and long-term recordings for the PEDOT and tungsten microelectrodes.

Fig. 3d presents the signal details of each state. The interictal state is characterized by a steady signal wave produced by physiological activity. In the preictal state, the signals are similar to interictal waveforms but gradually show allorhythmia. At this stage, epileptiform discharges consisting of spikes or sharp waves can sometimes be observed (Fig. S15). The postictal state is characterized by low-frequency and low-amplitude signal waves. Interestingly, the ictal state shows various trends with six kinds of characteristic waveforms. The ictal state generally begins with a slow oscillation wave (ⅰ) that successively changes to a fast wave (ⅱ) and hypsarrhythmia wave (ⅲ), with gradually increasing amplitude and frequency. The hypsarrhythmia wave (ⅲ) is characterized by spikes and a slow wave complex, which transitions to a spike wave. The spike wave (ⅳ) and following sharp wave (ⅴ) are the characteristic waveforms of the ictal state. The spike wave (ⅳ) is characterized by high amplitude and high frequency, while the sharp wave (ⅴ) is characterized by high amplitude and low frequency. The ictal state finally ends with a sharp wave with continuously decreasing amplitude (ⅵ) before transitioning to the postictal state. The recordings suggest that our PEDOT microelectrodes can sensitively capture multifarious changes in neural activities during epileptic seizures.

Time and frequency domain analyses were performed based on the intracerebral recordings of epileptic seizures. Fig. S16 shows the signal-to-noise ratio (SNR) comparison between the PEDOT and tungsten microelectrodes. The average signal amplitude of the PEDOT group was 1307.09 ± 431.33 μV, which was approximately 1.7 times larger than that (750.08 ± 124.85) of the tungsten group. The average noise levels of the two groups were 111.26 ± 31.81 μV and 98.50 ± 15.29 μV, respectively, and there was no significant difference between the two groups. As a consequence, the average SNR of the PEDOT group significantly increased from 7.6 ± 0.2 to 11.6 ± 0.8 compared with the tungsten group. Power spectrum density (PSD) analyses were also performed based on the epileptic signals. Similarly, the PEDOT group exhibited significantly higher power than the tungsten group in the frequency range associated with epileptic activity (0−30 Hz) (Figs. 3h and S17a).

Due to the favourable biocompatibility, the PEDOT microelectrodes exhibited better stability than the tungsten electrodes and enabled continuous intracerebral recordings for up to 8 weeks (12−15 seizures recorded in 6 mice for each group). The short-term (2 weeks after implantation) and long-term (6 weeks later) recordings of epileptic signals and their spectrograms are shown in Fig. S18. The average signal amplitudes and the average noise levels in the PEDOT group show no significant differences between the short-term and long-term recordings (Figs. 3e, f), presenting stable seizure signals with high SNRs (Fig. 3g). In contrast, the average signal amplitude in the tungsten group decreases by approximately 153.12 ± 64.91 μV (Fig. 3e), which leads to a significant difference in the SNRs between the short-term and long-term recordings (Fig. 3g). The PSD features of the epileptic signals also exhibit excellent stability in the PEDOT group, while they are noticeably reduced in the tungsten group (Figs. 3h and S17b, c).

### Neural decoding for seizure prediction

Epilepsy is a common neurological disorder characterized by uncertain seizures^63^. Without timely and appropriate treatment, epileptic seizures will cause loss of consciousness or perception and disorders of mood or other cognitive functions^64^. One key aspect of epilepsy care is seizure prediction, which involves decoding the abnormal neuronal activity occurring during the period before seizure onset to allow preventive action before seizure occurrence^65, 66^. Previous studies have mainly focused on the development of decoding algorithms and prediction models based on public electroencephalogram datasets^64^ and have paid less attention to the impact of neural interfacing techniques on decoding performance. As the proposed PEDOT microelectrodes can achieve intracerebral recordings with higher SNR and better stability than traditional tungsten electrodes, we anticipate that these microelectrodes can improve seizure prediction performance.

To effectively verify our hypothesis, seizure prediction was designed as a three-class problem. As illustrated in Fig. 4a, the raw intracerebral signals were first preprocessed by fast Fourier transforms as well as statistical and PSD analyses to extract features that were beneficial for neural decoding. Then, these features were input into the decoders to predict the epileptic states (interictal, preictal and ictal). We chose four traditional machine learning algorithms as the decoders for epileptic state classification, including the support vector machine (SVM), K nearest neighbour (KNN), decision tree (DT) and boosting algorithms, due to their simplicity and low computational costs in solving classification problems^67^. These four classifiers were first evaluated using the short-term recording signals. The classification accuracy was found to be essentially independent of the amounts of signal samples used for training (Fig. S20), suggesting the validity of the chosen classifiers. Although the classifiers show different accuracies depending on the algorithm complexity, the classification accuracies for the epileptic states are higher in the PEDOT group than in the tungsten group, with an average increase of approximately 2% in each case (Figs. 4b-e). The classification performance of each decoding algorithm was also assessed according to other statistical metrics, including the sensitivity (SEN), specificity (SPE), receiver operating characteristic (ROC) curve and area under the ROC curve (AUC) (Figs. S21a, S22a and Table S1). The results demonstrate higher SEN, SPE, and AUC values of the interictal and preictal classes in the PEDOT group than in the tungsten group. According to the signal analysis across different frequency bands, the improved classification performance can be attributed to the higher recording quality of neural signals in the PEDOT group, enabling better distinction between the interictal and preictal signals and thus more accurate seizure prediction (Fig. S19).

**Fig. 4.**
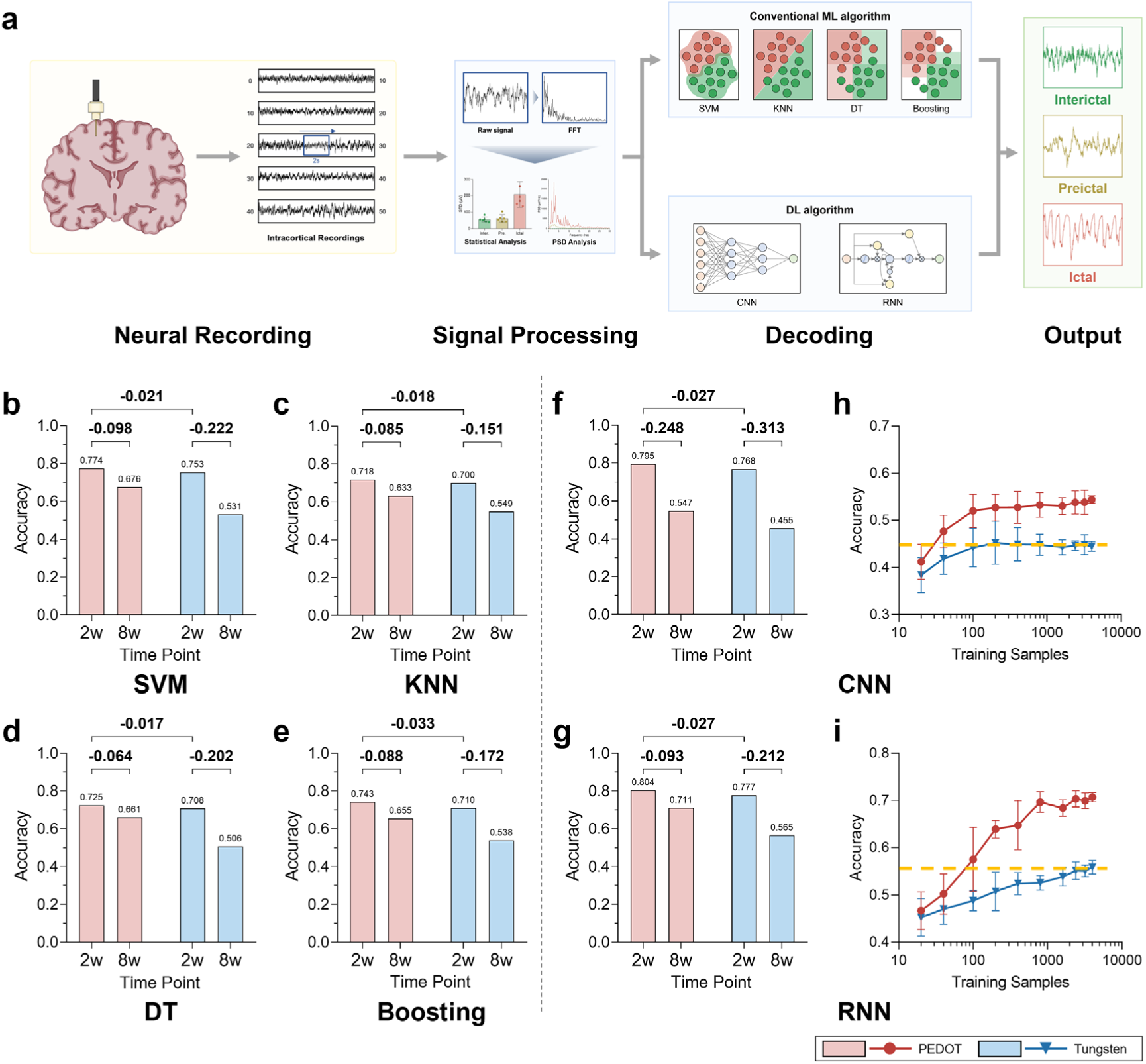
Neural decoding for seizure prediction. (**a**) A flow diagram of the decoding process and outputs of epileptic states. (**b**-**g**) Comparisons between the PEDOT and tungsten groups in terms of the classification accuracies of (**b**-**e**) four machine learning (ML) algorithms and (**f**, **g**) two deep learning (DL) algorithms based on the short-term (0−2 weeks) and long-term (6−8 weeks) recordings. (**h**, **i**) Classification accuracy as a function of the amount of training samples (mean ± STD) for the CNN and RNN algorithms, respectively.

In practical applications, the absence of chronic stable neural signals for metal-based microelectrodes remains a challenging problem for the durable decoding of intracerebral recordings. For example, in BCI applications, recording instabilities can deteriorate the decoding performance, making previously calibrated decoders no longer appropriate. To reveal the potential of our flexible PEDOT microelectrodes in stable decoding, long-term recordings collected 6−8 weeks after implantation were used as the testing set for the above four classifiers. As shown in Figs. 4b-e, all classifiers showed better performance using the long-term recordings in the PEDOT group than in the tungsten group. Compared with the performance using the short-term recordings, the classification accuracies were slightly decreased by 6−10% in the PEDOT group, while the accuracy was reduced by up to 15−22% in the tungsten group. This result is consistent with the more stable recoding performance of PEDOT microelectrodes. Similarly, the SEN, SPE, and AUC of each classifier in the PEDOT group were superior to those in the tungsten group (Figs. S21b, S22b and Table S2), confirming the improved decoding stability enabled by the flexible neural interfacing technique.

To further explore the influence of the proposed electrodes on decoding algorithms, we also used deep learning algorithms, which enables automatically learning of multilevel features. Two classic deep learning algorithms, namely, the convolutional neural network (CNN) and recurrent neural network (RNN)^68^, were chosen for seizure prediction. For the short-term recording dataset, the CNN and RNN algorithms both show robust accuracies of ∼0.8 (Figs. 4f-g) due to the high algorithm complexity. Moreover, the decoding stability was better in the PEDOT group than in the tungsten group, as shown by the better long-term classification accuracy (0.547 *vs.* 0.455 with the CNN and 0.711 *vs.* 0.565 with the RNN at 6−8 weeks postimplantation).

It has been well documented that the performance of deep learning algorithms strongly depends on the size of the dataset because the representation learning mode of deep learning requires sufficient training data to estimate the parameters^68^. Thus, we investigated how the amount of training data affects the decoding performance of deep learning algorithms in the PEDOT and tungsten groups. The classification performance of the CNN and RNN algorithms improved more significantly as the number of training samples increased in the PEDOT group than in the tungsten group (Figs. 4h, i) and plateaued when ∼400 training samples were used (0.544 ± 0.007 *vs.* 0.445 ± 0.009 with the CNN and 0.707 ± 0.009 *vs.* 0.559 ± 0.013 with the RNN, Figs. 5h, i). Owing to the rapid increase in the classification performance, the accuracies of the algorithms trained based on the PEDOT group quickly approached the maximal accuracy achieved in the tungsten group with considerably less training data, as indicated by the dotted line in Figs. 5h, i. This could particularly benefit the implementation of deep learning algorithms in clinical application scenarios, where acquiring large datasets is sometimes a difficult task due to the problem of data privacy^69, 70^.

### Warning system for epileptic seizures

Based on epileptic state decoding, we developed a warning system for seizure prediction. SVM algorithm with optimal simplicity and classification performance was used as the decoder. Similar to the previously reported warning system^71, 72^, we applied a decision threshold in our warning system, where an alarm was triggered if the number of preictal/ictal predictions for the incoming data in a defined period (*e.g.*, 20 seconds in our case) was higher than the decision threshold. This threshold ensures that the seizure alarms were triggered only when several preictal/ictal predictions were in close temporal proximity.

When the threshold was more than 7, namely, > 7 preictal/ictal predictions in 20 seconds, the number of false alarms was effectively reduced in our system (Fig. S23). Therefore, a = 7 threshold was used to evaluate the performance of our warning system. Figs. 5a-b displays an example of our proposed seizure alarm system based on the segments containing whole epileptic seizure events recorded by the PEDOT and tungsten microelectrodes, respectively. When the threshold was set to 8, there were more false alarms in the interictal state for the tungsten group than for the PEDOT group, and desirable alarms in the preictal/ictal state occurred more frequently for the PEDOT group (Figs. 5a-b).

**Fig. 5.**
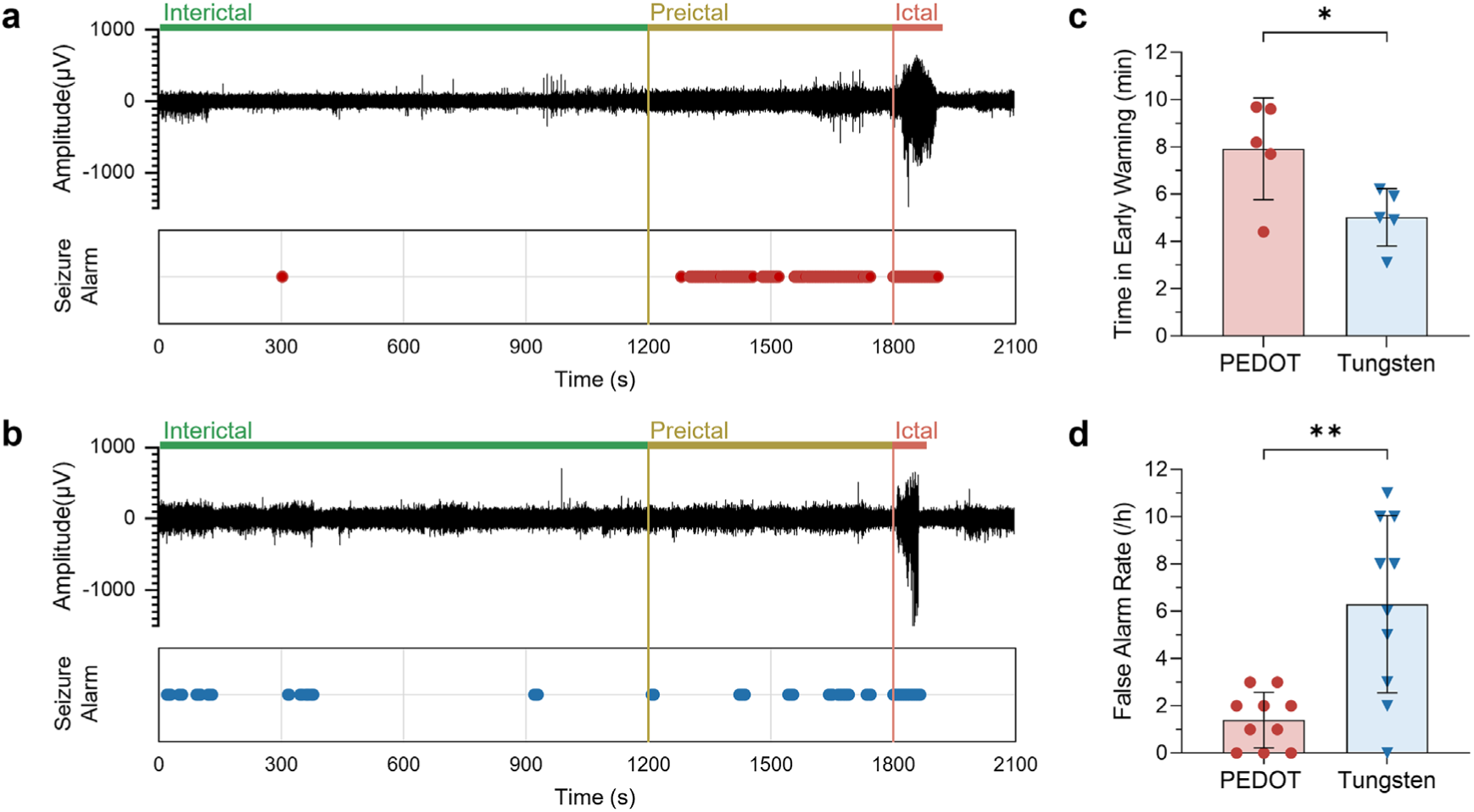
Alarm system for epileptic seizures. (**a**, **b**) The applied examples of the seizure alarm system using recordings from the PEDOT (**a**) and tungsten (**b**) groups. The ictal state (seizure) begins at 1800 seconds, and the period of 600 seconds (10 minutes) before seizure onset is defined as the preictal state. (**c**) Early warning time of the two groups at the threshold of more than 7 (*n* = 5 for each; student’s independent-samples t tests, *t* = 2.627, *p* < 0.05). (**d**) False alarm rates of the two groups at the threshold of more than 8 (*n* = 10 for each; student’s independent-samples t tests, *t* = −3.950, *p* < 0.01).

We quantitatively compared the performance of two groups by calculating two clinically relevant metrics, namely the false alarm rate (the average number of false alarms triggered per hour during the interictal period) and the early warning time (the average alarm duration before seizure onset), which were introduced in previous studies^71, 73–75^. The PEDOT group demonstrated an early warning time of 8 ± 2 minutes before seizure onset, which is earlier than that of the tungsten group (5 ± 1 minutes), as shown in Fig. 5c. This extended period provides patients and healthcare professionals with more time to take necessary actions, such as neuromodulation or drug delivery, in a closed-loop manner to mitigate incoming seizures. Furthermore, the PEDOT group had a lower false alarm rate of 1.4 ± 1.1 per hour compared to the false alarm rate of 6.3 ± 3.6 per hour in the tungsten group (Fig. 5d). This lower false alarm rate minimizes unnecessary disturbance from alarms in patients’ daily lives. The optimal warning performance of the PEDOT group is consistent with its improved decoding performance. As exemplified in Video 3, we have successfully established a real-time seizure warning system through decoding the neural signals recorded by both PEDOT and tungsten microelectrodes, demonstrating the promising clinical utility of flexible electronics in the prediction of seizures.

## Discussion

In this work, we used PEDOT microfibers as flexible neural electrodes to investigate how flexible neural interface techniques can be combined with machine learning algorithms to enable long-term reliable neural decoding performance. We synthesized conductive polymer microfiber electrodes with desirable mechanical, electrochemical and biocompatible properties for neural recordings, assessed their interplay with various machine learning algorithms and established a real-time seizure warning system. Thus, our research provides a typical example of integrating flexible implanted electronics with artificial intelligence analysis to overcome the intrinsic trade-off between accuracy and stability in neural decoding based on conventional rigid electrodes.

In the last decade, flexible electronics have emerged as the next-generation neural interfacing technique to address the limitations of conventional rigid electronics. Despite the significant progress made on the front-end integration of flexible electronics with the brain for long-term recordings, limited attention has been given to back-end input/output connectivity and data processing, which is a key step to provide closed-loop feedback for neural interfacing devices^76^. Our work fills the existing gap between signal recording and processing with flexible neural electronics and, for the first time, demonstrates their unique capability in improving the longitudinal accuracy of neural decoding and providing long-term treatment of chronic neurological disorders such as epilepsy. This could promote research on flexible neural electronics from fundamental neuroscience to practical applications such as precise electronic medicine^76^.

Our work reveals the great potential of exploring flexible BCIs^77^. The performance of conventional BCI systems based on rigid electronics declines over time due to instabilities in the neural recording. Flexible electronics address this issue and could offer a universal strategy for stabilizing BCI performance based on reliable neural interfaces, which is different from existing methods focused on improving the algorithms. Furthermore, the use of flexible neural electronics enables good decoding performance in scenarios with less training data. This could improve the clinical viability of BCIs by reducing the large amount of patient-specific healthcare data required for accurate decoding and reducing the training time and computational cost of decoders.

In the future, there is ample room for further integration of flexible electronics and mechanical learning in neural decoding. Considering the processability and multifunctionality of PEDOT fibers, they can be assembled into fiber microelectrode arrays or combined with various neuroimaging techniques (*e.g.*, MRI) to collect multimodal information on neural activity in a scalable manner. This can be used to build crossover models to improve the decoding performance in broad applications. Moreover, flexible neural recoding electrodes can be integrated with other flexible devices with data transmission and processing functions (*e.g.*, wireless streaming^78^, stretchable neuromorphic chips^79^) to achieve flexible BCIs.

## Material and methods

### Fabrication and electrochemical characterization of the PEDOT fibers

The PEDOT fibers were prepared through a previously developed wet spinning technique^47–49^. PEDOT:PSS aqueous dispersions with 1.3 *wt*% and 3.0 *wt*% were used as the spinning dopes. The low-concentration dispersion was purchased from Heraeus (Clevios^TM^ PH1000), and the high-concentration dispersion was prepared by rotary evaporation from the initial dispersions. The dopes were contained in a 10 mL plastic injector (14.9 mm inner diameter) and mounted on a syringe pump (LSP01-2A, Hangzhou Noted). Then, they were injected into a coagulation bath (98 *wt*% sulfuric acid) through a stainless-steel needle (21−27 G) at an injection speed of 4 mL min^−1^ at 25 °C. Note that the dopes should be injected into the bottom of the sulfuric acid solutions. With this approach, the PEDOT:PSS dopes with densities lower than that of sulfuric acid float in the medium and quickly solidify into hydrogel fibers, which were continuously collected on a wheel. The as-obtained hydrogel fibers were immersed in sulfuric acid for one hour and washed with water 3−5 times. Finally, the hydrogel fibers were hung to dry at 50 °C for 24 h, and the ends of the fibers were tied to 10 g weights. For comparison, the hydrogel fibers were also hung without the weights.

The electrochemical properties of the dried PEDOT fibers were measured on a commercial electrochemical station (CHI660e, Shanghai Chenhua) through a three-electrode setup, where an Ag/AgCl electrode and Pt gauze were used as the reference electrode and counter electrode, respectively. One end of a PEDOT fiber (∼4 mm length) was sticked to Pt wire by conductive silver paint, and the other end was immersed into 10 mM PBS. The immersion depth was controlled to be ∼1-2 mm to avoid the contact of the Pt wire with the PBS electrolyte. The impedance spectra were acquired with a 5-mV voltage amplitude sine wave in the frequency range of 10^−1^−10^5^ Hz. The charge storage capacity was estimated through cyclic voltammetry tests in the potential window of 0.5 V at scan rates of 10−200 mV s^−1^. The charge injection capacity was assessed through bipolar pulse voltage stimulation under a voltage amplitude of 0.5 V and different pulse periods from 100 μs to 10 ms.

### Experimental animals and seizure induction

CR (CD-1) male mice (8−9 weeks) were used and purchased from Weitong Lihua Experimental Animal Technology Co., Ltd. (Zhejiang Province, China; animal certificate number: SCXK (Zhejiang) 2019–0001). The mice were housed in individual cages on a 12-hour light-dark cycle with controlled temperature (21−25 °C), humidity (40−60%) and ad libitum access to chow and water. All procedures to maintain and use the mice were reviewed and approved by the Laboratory Animal Ethics Committee of Jinan University, China (IACUC-20220621-07). In this study, temporal lobe epilepsy (TLE) was induced via pilocarpine, which allowed us to acquire spontaneous seizure models with high efficiency. Briefly, after a quarantine period of 10 days, the mice were intraperitoneally injected with 1 mg kg^−1^ scopolamine (1 mg mL^−1^, dissolved in ultrapure water), followed by injection with 300 mg kg^−1^ pilocarpine (150 mg mL^−1^, dissolved in ultrapure water) half an hour later. Diazepam (5 mg mL^−1^) was used to terminate acute seizures 1 hour after the first seizure.

### Fabrication and surgical implantation of neural microelectrodes

For intracerebral recording experiments, the PEDOT fibers were insulated with a 2 μm thick layer of Parylene-C through a vacuum vapour deposition process, with the cross sections exposed as electrically active sites via mechanical cutting. One end of the single-filament PEDOT fibers was then attached to the connecting wire with conductive silver paint, thereby assembling the implantable neural microelectrodes. The tungsten microelectrodes were constructed using a similar method.

The mice were anaesthetized with isoflurane (induction 5%, maintenance 2−3%; Sigma-Aldrich, USA) and secured in a stereotaxic instrument (RWD, China). Following a midline incision, a small craniotomy and microelectrode implantation were performed at the ventral hippocampus (AP: −2.3 mm; ML: −1.5 mm; DV: −1.6 mm from the skull). As the PEDOT microelectrodes were too flexible for self-supported intracerebral implantation, a conventional drawing lithography method for temporary maltose coating^80^ was used to assist the implantation process (Figs. S11a-c). Brass screws, which were used as the reference and ground electrodes and to support the dental cement, were severally fixed to the skull away from the implantation sites. To keep the trauma of the implantation site consistent, the tungsten microelectrodes were implanted in the same manner. After cleaning the surface of the skull with hydrogen peroxide, the microelectrodes were secured to the skull although light-curable dental cement (RIVA®, SDI, Australia). During surgery, a heat pad was applied to maintain the body temperature at 37 °C, and erythromycin eye ointment was used to prevent corneal drying.

### Intracerebral recordings and analysis

A CED 1401 acquisition board (Cambridge Electronic Design, UK) and Spike2 software (Version 8.0, Cambridge Electronic Design, UK) were used to collect intracerebral neural signals in freely moving mice at a sampling rate of 500 Hz. Note that there was constant 50 Hz harmonic noise, which was due to electromagnetic interference from the mains regardless of proper grounding or shielding. This noise was removed using a built-in script (HumRemoveExpress, https://ced.co.uk/us/tutorials/spike2datahr) in the Spike2 software in the following signal analyses. A digital video camera was used to simultaneously monitor behaviour during the recordings. The short-term recordings of epileptic seizures were carried out during the first 2 weeks after microelectrode implantation, and the long-term recordings were completed 6 weeks later.

Time and frequency domain analyses were performed using MATLAB software (Version R2020b, MathWorks, USA). To compare the signal-to-noise ratios (SNRs) of the two microelectrodes, the signal amplitude was calculated as the average peak-to-peak amplitude of the characteristic spike and sharp wave during a seizure, and the noise level was defined as two times the standard deviation of that during awake recordings between seizures. Spectrograms were converted from raw signals using short-time Fourier transform (STFT). The power spectrum density (PSD) was calculated in the frequency domain using Welch’s method. Different frequency bands, including the delta band (0.5−4 Hz), theta band (4−8 Hz), alpha band (8−13 Hz), and beta band (13−30 Hz), were acquired by finite impulse response (FIR) filters, and the average power across frequency bands was calculated using the bandpower function in MATLAB. In the above analysis, the parameters were as follows: FFT length = 1024; window length = 1024; overlap = 512; sampling rate = 500 Hz.

### Magnetic resonance imaging

MRI scanning was performed in a 9.4 T MR scanner (Bruker BioSpin MRI, Germany) with an 86 mm volume coil for transmission and a 2-cm diameter single-loop surface coil for receiving (ParaVision Version 6.0.1 for MRI acquisitions). Mice were anaesthetized with 4% isoflurane, followed by 2% isoflurane delivered via a nose cone to maintain anaesthesia during MRI scanning. The body temperature, heart rate, and respiration were all monitored and within normal ranges. The MRI surface receiver coil was fixed over the mouse heads, and the PEDOT fiber microelectrodes passed through this coil. A T2-weighted coronal plane scan was performed after the calibration scan. The parameters were as follows: TE = 33 ms, TR = 2500 ms, slice thickness = 0.8 mm, matrix size = 256 × 256, and field of view = 20 × 20 cm².

### Immunohistochemistry staining

The anaesthetized mice were transcardially perfused with 0.9% saline followed by 4% paraformaldehyde (PFA; Solarbio, China) in 0.1 m phosphate-buffered saline (PBS). The brains were removed and postfixed with 4% PFA overnight at 4 °C and then transferred to 30% sucrose at refrigerated temperature to aid in cryoprotection. The postfixed brains were frozen in Tissue Tek OCT at −80 °C and sliced coronally into 20 μm sections with a cryostat machine (CM1900, Leica Microsystems, USA). The sections were washed with 0.1 m PBS three times for 15 min each and blocked for 1 hour with a solution of 10% normal goat serum (Vector Laboratories, USA) and 0.3% Triton X-100 (T8787, Sigma-Aldrich, USA) in 0.1 M PBS. Then, the sections were immunostained by incubating for 2 days at 4 °C with the appropriate primary antibodies, including rabbit anti-ionized calcium binding adaptor molecule 1 (Iba1) for microglia (1:100, ab289370, Abcam, UK), mouse anti-glial fibrillary acidic protein (GFAP) for astrocytes (1:500, ab279290, Abcam, UK), rabbit anti-neuronal nuclei (NeuN) for neuronal densities (conjugated with Alexa Fluor® 647, 1:1000, ab190565, Abcam, UK), and rabbit anti-Aquaporin 4 (AQP4) for aquaporin (1:500, ab259318, Abcam, UK). After six rinses in 0.1 M PBS, the sections were incubated for 6 hours at room temperature with DAPI (1:1000, Sigma-Aldrich, USA) and the following secondary antibodies: goat anti-rabbit Alexa Fluor® 555 (1:200, ab150078, Abcam, UK) and donkey anti-mouse Alexa Fluor® 488 (1:200, ab150105,

Abcam, UK). Finally, the stained sections were washed and mounted on glass slides with coverslips and imaged with ZEN microscopy software (Version 3.6, Carl Zeiss GmbH, Germany) on a confocal laser scanning microscope (LSM 880, Carl Zeiss GmbH, Germany).

### Whole-cell patch-clamp recordings

The brains were extracted from the anaesthetized mice and cut into 300 μm thick horizontal slices with a fully automated vibrating blade microtome (VT1200 S, Leica, Germany) in freezing cutting solution (containing 85 mM NaCl, 4 mM MgCl_2_, 2.5 mM KCl, 0.5 mM CaCl_2_, 24 mM NaHCO_3_, 1.25 mM NaH_2_PO_4_, 75 mM sucrose, and 25 mM glucose). The brain slices were incubated in oxygenated *N*-methyl-d-glucamine (NMDG) artificial cerebrospinal fluid (NMDG-ACSF) (containing 93 mM NMDG, 20 mM HEPES, 2.5 mM KCl, 0.5 mM CaCl_2_, 30 mM NaHCO_3_, 1.2 mM NaH_2_PO_4_, 10 mM MgSO_4_⋅7H_2_O, 5 mM sodium ascorbate, 3 mM sodium pyruvate, 2 mM thiourea, and 25 mM glucose; pH 7.3 adjusted with HCl; 300∼310 mOsm/L) at 34 °C for 30 min and then maintained at room temperature. Electrophysiological recordings were performed with a patch-clamp amplifier (Axon MultiClamp 700B, Molecular Devices, CA) in oxygenated ACSF (containing 126 mM NaCl, 2 mM MgCl_2_, 2.5 mM KCl, 2 mM CaCl_2_, 26 mM NaHCO_3_, 1.25 mM NaH_2_PO_4_, and 10 mM glucose; pH 7.3 adjusted with NaOH; 310∼320 mOsm/L). Patch-clamp pipettes were fabricated from stretched borosilicate glass (approximately 5−8 MΩ) and filled with a pipette solution (containing 110 mM HEPES, 4 mM KCl, 0.3 mM Na_2_GTP, 4 mM Mg_2_ATP, 26 mM K-Gluconate, 10 mM Phosphocreatine; pH 7.3 adjusted with KOH; 290 mOsm/L). Electrophysiological data were collected and analysed with pClamp software (Version 10, Molecular Devices, CA) and Clampex software (Version 10.5, Molecular Devices, CA). The data were sampled at 10 kHz and filtered at 3 kHz.

### Machine learning for neural decoding

The available neural decoding data included interictal, preictal, and ictal signals at a ratio of 10:5:1. These signals were identified and labelled by neurological experts. Since there is clearly an overabundance of available interictal and preictal data, it is typical to select such an imbalanced ratio. In addition, there is no consensus on the exact length of abnormal activation before seizure occurrence, which is typically between 10 and 120 minutes. In this study, the preictal signals were defined as the segment 10 minutes before a seizure period. For epileptic signal decoding, four traditional machine learning algorithms, namely, the support vector machine (SVM), K nearest neighbour (KNN), decision tree (DT), and boosting algorithms, as well as two deep learning algorithms, namely, the convolutional neural network (CNN) and recurrent neural network (RNN) algorithms, were selected as the classifiers. Seventy percent of the short-term recording data were used as the training set, and the remaining 30% were used as the testing set, while the long-term recordings were used only as the testing set. Before training and testing, the raw signals were digitally filtered between 0.5 Hz and 30 Hz, and divided into frames with lengths of 2 seconds and an overlap of 1 second. The time domain (for instance, statistical analysis with standard deviation and root mean square error) and frequency domain (including fast Fourier transforms and PSD analyses) were both used for feature extraction. For a fair comparison, all processing parameters and settings were the same in both groups. The code to process and analyse neural data in this study is available at https://github.com/Huangzx1023/Machine-learning-for-epileptic-state-classification.

All decoding algorithms were replicated at least 3 times, and one of the results was chosen as a detailed demonstration for each algorithm. A confusion matrix was used as a systematic approach to illustrate the classification accuracy of each algorithm. The statistical measures for comparing the decoding performance between groups included the most widely used metrics, including the accuracy (ACC), sensitivity (SEN), and specificity (SPE), which were calculated as follows:

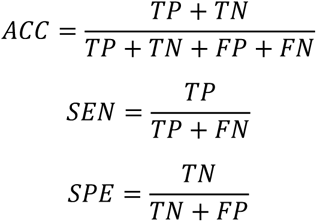

True positive (TP) denotes that the positive class was predicted correctly; true negative (TN) denotes that the negative class was predicted correctly; false positive (FP) denotes that the negative class was incorrectly predicted as positive; and false negative (FN) denotes that the positive class was incorrectly predicted as negative. Furthermore, the receiver operating characteristic (ROC) curve was used to assess the true positive and false positive rates between two groups, and the area under the ROC curve (AUC) was applied to quantify the performance for each class.

### Seizure warning system design

The trained classifier with the best performance was chosen as the underlying algorithm in the proposed system. Continuous signals from 2-second windows were used as inputs to predict epileptic states. A decision threshold (from 0 to 9) was set for the number of preictal or ictal predictions made in the past 20 seconds. Alarms would be triggered for seizure, once the number of preictal/ictal predictions crossed the decision threshold. The period 10 minutes before seizure onset was defined as the preictal period, where the triggered alarms were defined as true alarms. The period that was neither preictal nor ictal was defined as the interictal period, and the number of alarms triggered per hour in this period was used to determine the false alarm rate. The time that an intensive alarm occurred within the 10-minute period before a seizure was defined as the beginning of early warning, and the alarm duration was counted as the early warning time.

### Statistical analysis

Statistical comparisons were performed using SPSS (Version 25.0, SPSS Inc., USA) and GraphPad Prism software (Version 8.0.1, GraphPad Software Inc., USA). Statistical significance was tested with student’s independent-samples or paired-sample t tests and DeLong’s test. The equality of the variance was tested with Levene’s test. All data are presented as the mean ± STD. The significance level was indicated using * for *p* < 0.05, ** for *p* < 0.01, *** for *p* < 0.001 and *ns* for no significant difference. The error bars represent the STD.

## Supporting information

Video 1. Flexible PEDOT fiber for bulb lighting test

Video 2. An example for intracerebral recordings of epileptic seizure

Video 3. Examples for real-time seizure warning system

## Acknowledgement

This work is finacially supported by the National Natural Science Foundation of China (82227802, 81901709, 81871323, 32200796) and the Science and Technology Planning Project of Guangdong Province (2019A1515011918).

## Author Contributions

Z.Y. X. conceived the study and designed the experiments with the help form Z.C.H. and S.X.Z. Z.Y. X. fabricated and characterized the PEDOT microfibers and Z.C.H. fabricated the neural microelectrodes. Z.C.H. and J.Z. conducted the biological and tissue experiments with assistance from S.W. and Q.H. J.W.D. and Z.X.H. wrote the code to process and analyze the neural data. Z.C.H. and Z.J. analyzed the rest of data. Z.C.H. and Z.Y.X. wrote the manuscript with advices from K.F.S. and S.X.Z. Z.Y.X. and S.X.Z. supervised the study. All authors contributed to the article and approved the submitted version.

## Ethics declarations

All procedures to maintain and use the mice were reviewed and approved by the Laboratory Animal Ethics Committee of Jinan University, China (IACUC-20220621-07).

