## Supporting Information for "Long-term reliable neural decoding based on flexible implantable microelectronics and machine learning for seizure prediction application"

### for

#### Section 1. Supplementary experimental procedures

##### S1.1 Materials and Synthesis

The Ag/AgCl electrode and Pt wire (500 μm diameter) were purchased from Shanghai Chenhua. The Pt gauze (100 mesh) was brough from Sigma Aldrich (product number, 298093). Tungsten wires (80 μm diameter) were on-line purchased from JingDing Metal in Alibaba platform. Deionised water with resistivity >18 MΩ cm was prepared by a Milli-Q system and used for all experiments. The conductive silver paint was obtained from SPI supplies. The chemicals used in this work were bought from Sigma Aldrich unless otherwise specified.

##### S1.2 Characterisation of PEDOT fibers

The morphology of PEDOT fibers was characterised by a field-effect scanning electron microscope (SEM, Hitachi SU8010, Shimadzu Corporation) at 5 kV voltage. Wide-angle X-ray scattering (WAXS) measurement were conduted in the Xenocs Xeuss 2.0 SAXS/WAXS system in transmission mode. The source was GeniX3D Cu ULD 8 keV with wavelength of 1.54 Å. Generally, 3~5 PEDOT fibers were aligned into a bundle and placed on an aperture card. The aperture card was then transferred to the WAXS sample holder and placed at 101.17 mm from the 2D detector (Dectris Pilatus 200 K). The exposure time was 600 s. Data processing was performed using the software Foxtrot provided by Xenocs. The Herman's orientation factor ($f$) was calculated by the equation to estimate the orientation degree of PEDOT microcrystals[1,2]:

$$f = \frac{3\,(\cos\theta)^2 - 1}{2}$$

where the mean-square cosine was calculated from the scattered intensity $I(\theta)$ by integrating over the azimuthal angle ($\theta$) through the equation[1,2]:

$$(\cos\theta)^2 = \frac{\int_0^{\pi/2} I(\theta)\sin\theta\,(\cos\theta)^2\,\mathrm{d}\theta}{\int_0^{\pi/2} I(\theta)\sin\theta\,\mathrm{d}\theta}$$

The electrical conductivit was measured on a electrochemical station using a two-probe method. Cyclic voltammetry (CV) test was performed to measure the resistance ($R$) of PEDOT fiber at a scan rate of 10 mV $s^{-1}$ within the potential window of -0.1~0.1 V. The electrical conductivity ($\sigma$) was extracted according to the equation:

$$\sigma = \frac{l}{R * S}$$

where $l$ and $S$ are the length and lateral area of the PEDOT fibers, respectively. The uniaxial tensile tests of the PEDOT fibers were conducted on a commercial testing machine (Instron 5943) at ambient temperature. The fiber length between jaws was set as 1 cm. The tensile rates for wet and dried PEDOT fibers were set as 15% strain per minute and 100% strain per minute, respectively. The tensile strength ($\sigma_b$) of the fibers was calculated by the equation:

$$\sigma_b = \frac{F}{S}$$

where F is the fracture stress. For the calculation of electrical conductivity and tensile strength of the fibers, their lateral area was estimated from the cross-sectional SEM iamges through ImageJ software considering the irregular cross-sectional shape. Typically, three fiber samples were used in measuring the electrical conductivity and tensile strength.

**S1.3 Electrochemical performance of PEDOT fibers and tungsten wires**

The charge storage capacity (CSC) and charge injection capacity (CIC) were obtained by integrating the cathodal part of $i{\sim}V$ curves in CV measurement and the cathodal part of $i{\sim}t$ curves in bipolar pulse voltage stimulation test, respectively. The following is the equations for the calculation of CSC and CIC[3,4]:

$$CSC = \frac{\int_{V_1}^{V_2} idV}{S * v}$$

$$CIC = \frac{\int_{t_1}^{t_2} i\,dt}{S * V}$$

where $v$ is the scan rate, $S$ is the immersed area of the electrodes, $V_1$ and $V_2$ are the initial and terminal points of the voltage window in the CV test; $t_1$ and $t_2$ are the initial and terminal points of pulse duration and $V$ is the amplitude of the applied voltage in the stimulation test.

**S1.4 Calculation of the bending stiffness PEDOT fibers and tungsten wires**

The geometrical and mechanical properties of PEDOT microfiber and tungsten wire used for the calculation is listed below.

| Materials | Cross section | | Young's modulus (GPa) | Poisson's ratio |
|---|---|---|---|---|
| | shape | dimension | | |
| PEDOT microfiber | rectangular | ~70 μm width<br>~20 μm thickness | ~9 | ~0.3 |
| tungsten wire | round | 80 μm diameter | 411 | 0.28 |

As the cross section of PEDOT microfiber is rectangular, its bending stiffness along the thickness direction is roughly estimated using the equation for the calculation of the bending stiffness ($J$) of an isotropic membrane[4]:

$$J = \frac{E * w * t^3}{12\,(1-\sigma)}$$

where $E, w, t$ and $\sigma$ are the Young's modulus, width, thickness and Poisson's ratio of the membrane, respectively. Therefore, the bending stiffness of PEDOT microfiber is calculated to be $6.0\times 10^{-10}$ N $\mathrm{m}^2$.

On the other hand, the bending stiffness of tungsten wire with round cross section can be calculated by the following equation[5]:

$$J = \frac{\pi * E * d^4}{64\,(1-\sigma)}$$

where $d$ is the diameter of tungsten wire. Therefore, the bending stiffness of tungsten wire is calculated to be $1.2\times 10^{-6}$ N $\mathrm{m}^2$, which is 3~4 orders of magnitude higher than the one of PEDOT microfiber.

Note that the shape of the cross section of PEDOT microfiber is not perfectly rectangular. The above values are only estimated values for comparison.

#### Section 2. Supplementary figures and tables

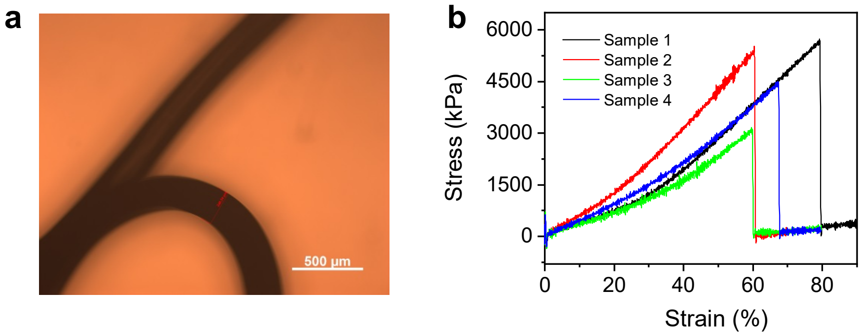


**Figure S1**. (a) Optical microscopic image and (b) Tensile stress-strain curves of hydrogel fibers spun from PEDOT:PSS dispersions. The hydrogel fibers are highly elastic a tensile strength of ~5 MPa and a large breaking strain of 50−80%, thereby facilitating the drying of PEDOT hydrogel fibers under drawing stress. The high elasticity could be related with the formation of numerous PEDOT microcrystal regions embedded in the network of PSS chain, as supported by the WAXS patterns in Figure S3.

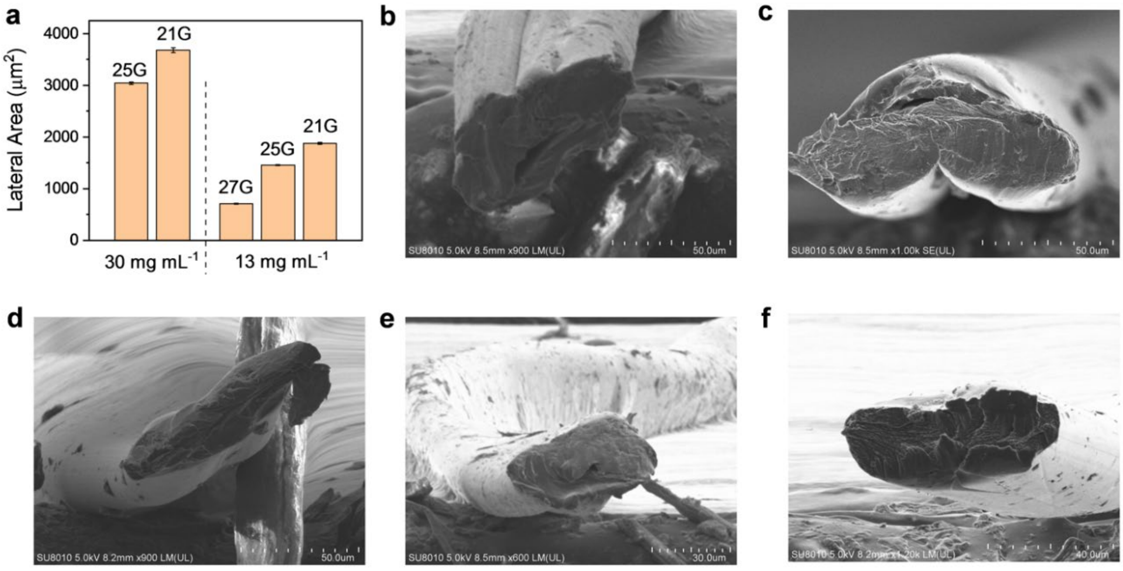


**Figure S2**. (a) Lateral area of the PEDOT fibers prepared under different conditions, including concentrations of spinning dopes and diameters of needles. (b-f) Scanning electron microscope images of the cross section of PEDOT fibers. The spinning conditions are (b) 30 mg $mL^{-1}$ PEDOT:PSS dispersion, 25G needle, (c) 30 mg $mL^{-1}$ PEDOT:PSS dispersion, 21G needle, (d)

13 mg mL$^{-1}$ PEDOT:PSS dispersion, 21 G needle, (e) 13 mg mL$^{-1}$ PEDOT:PSS dispersion, 27G needle, and (f) 13 mg mL$^{-1}$ PEDOT:PSS dispersion, 25G needle.

The lateral dimension of the PEDOT fibers is found to increase with the concentration of PEDOT:PSS dispersions and the inner diameter of the needles (Figure S2a). Moreover, the PEDOT fibers show an irregular cross section instead of round shape (Figure S2b-f). In most cases, ribbon-like shape can be observed. This could be a result of the asymmetrical collapse of the injected PEDOT:PSS dopes during the ultrafast dehydration process induced by concentrated $H_2SO_4$. As seen from the calculation in S1.4, the ribbon-like shape greatly contributes to the flexibility of PEDOT fibers.

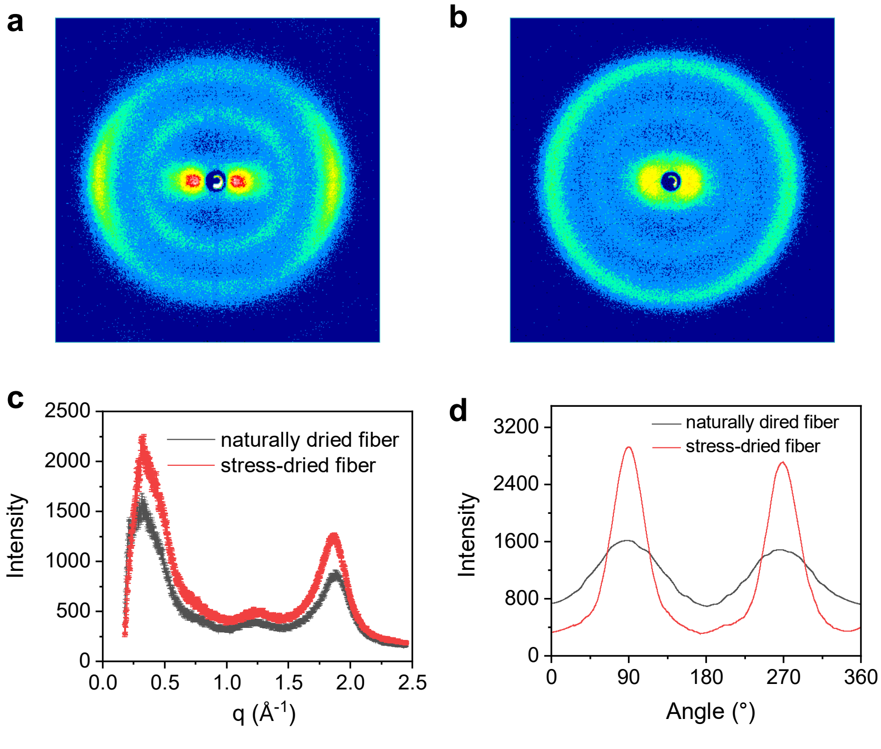


**Figure S3**. (a-b) The two-dimensional (2D) WAXS patterns of PEDOT fiber prepared through (a) stress drying and (b) natural drying. (c) The X-ray scattered intensity versus the scattering vector and (d) Angular scattering intensity profile for naturally dried and stress-dried PEDOT fibers.

The crystalline structure of the PEDOT fibers was characterized by WAXS. The characteristic arcs in the 2D scattering patterns of stress-dried fibers indicates preferred orientation of the PEDOT crystals along the fiber axis (Figure S3a). As contrast, no crystalline orientation is found in the naturally dried fibers, as indicated from the homogeneous intensity of the scattering rings (Figure S3b). The strong scattering at $q$ = ~1.8 $Å^{-1}$ should be attributed to the π-π lamella stacking of PEDOT crystals[1,2]. It is found that the stress-dried PEDOT fibers have higher scattering intensity at $q$ = ~1.8 $Å^{-1}$ than the one of naturally dried fibers (Figure S3c), indicating higher crystallinity for stress-dried fibers. Furthermore, the angular scattering intensity profile of stress-dried PEDOT fibers is narrower (Figure S3d); the calculated Herman's orientation factor of stress-dried fibers is also higher than the one of naturally dried fibers (0.71 vs 0.12), supporting the enhanced orientation of PEDOT crystals under the stress.

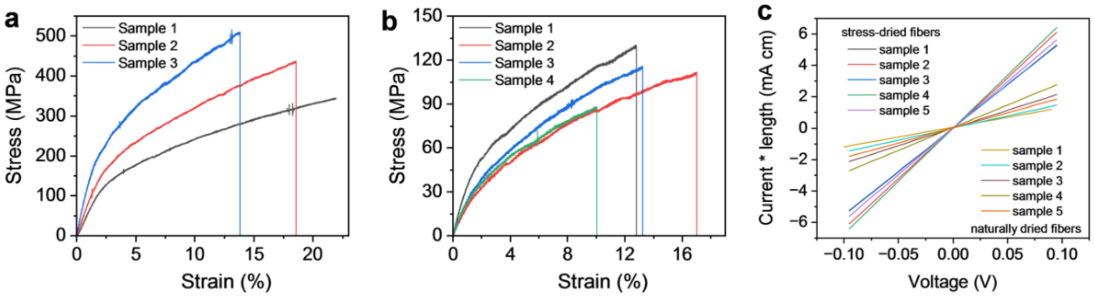


**Figure S4**. (a-b) Tensile stress-strain curves for (a) stress-dried PEDOT fibers and (b) naturally dried PEDOT fibers. (c) Current-voltage curves for stress-dried and naturally dried PEDOT fibers in two-electrode resistance measurement.

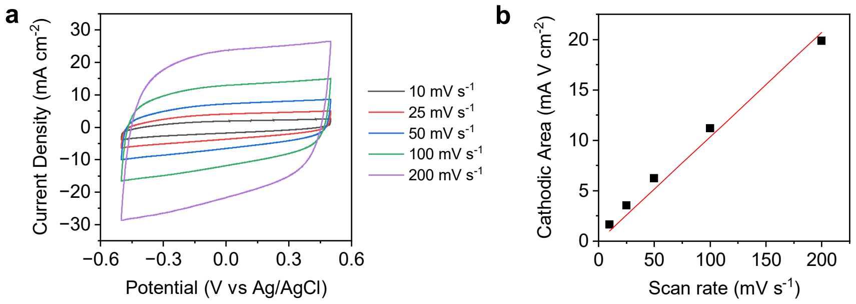


**Figure S5**. (a) Current-potential curves and (b) the cathodic area of PEDOT fibers in cyclic voltammetry tests under different potential scan rates.

The CV curves of PEDOT fibers in PBS is nearly rectangular under wide-ranged potential scan rates of 10−200 mV $s^{-1}$. Meanwhile, a linear relationship between the scan rate and the cathodic area in the CV curves can be observed. These results suggest facilitated ion storage in PEDOT fibers, enabling low impedance as shown in Figure 1f.

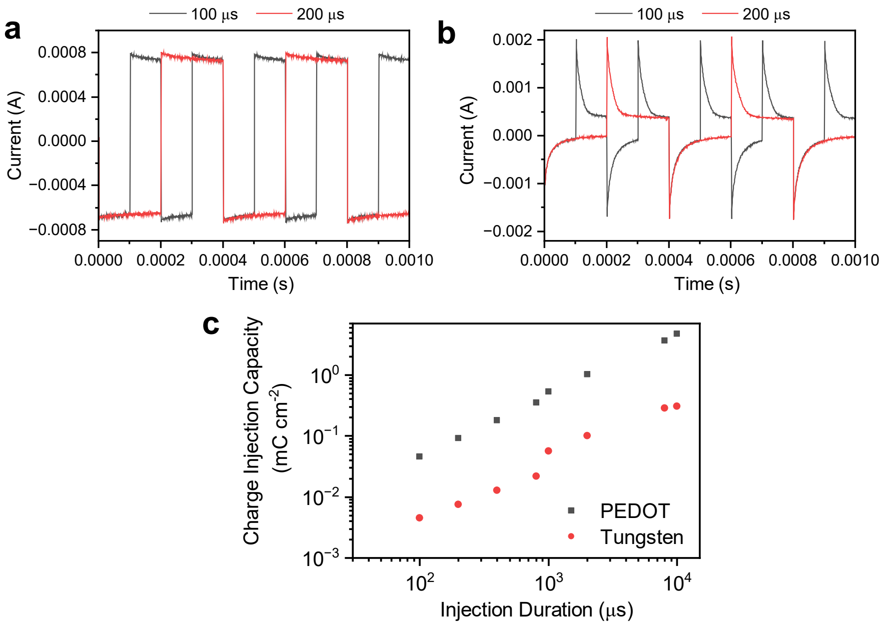


**Figure S6**. (a-b) Current-time curves of (a) PEDOT fiber and (b) tungsten wires in bipolar pulse voltage stimulation measurement. The voltage amplitude is 0.5 V. The pulse duration for charge injection is 100 and 200 μs. The decrease of the current is slow for PEDOT fibers after pulse voltage stimulation owing to the large interfacial capacitance[6]. As contrast, the current reduces quickly for tungsten wires due to the small electrical double layer capacitance. (c) The charge injection capacity of PEDOT fiber and tungsten wires under different pulse durations for charge injection.

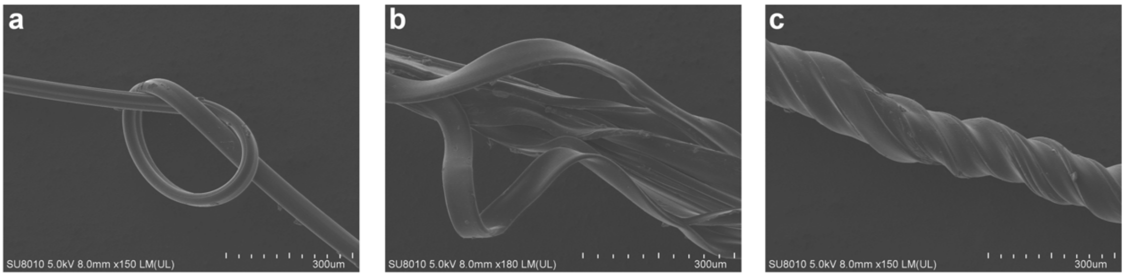


**Figure S7**. SEM images of PEDOT fibers after (a) knotting and (b-c) twisting. The knotting and twisting tests suggest the good mechanical flexibility and toughness of PEDOT fibers.

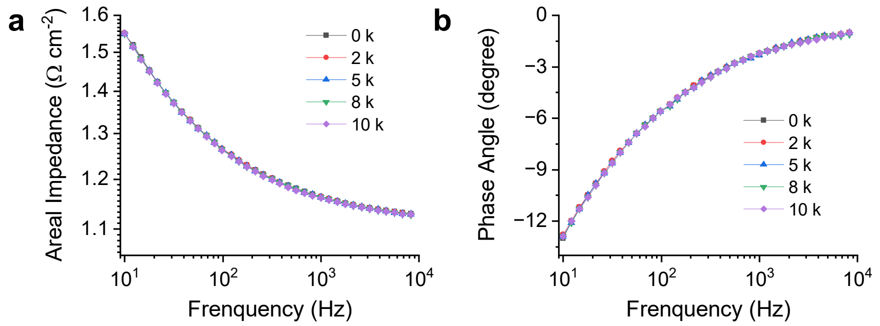


**Figure S8**. (a-b) The change of (a) impedance and (b) phase angle of PEDOT fibers with frequency after different times (0~10000) of bending cycles at 180° bending angles. There is almost no change for the impedance and phase angle of the PEDOT fibers after 10000 bending cycles, indicating good mechanical stability for use in neural recording.

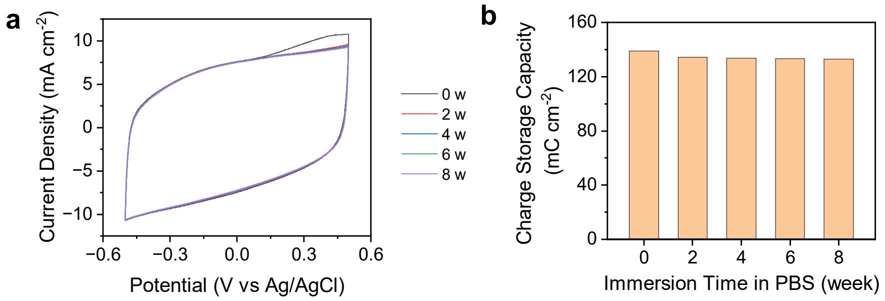


**Figure S9**. (a) Cyclic voltametric curves of PEDOT fibers at the potential scanning rate of 50 mV $s^{-1}$ after different immersion times in PBS from 0 week to 8 weeks. (b) The change of charge storage capacity of PEDOT fibers with the immersion time. The CSC of PEDOT fiber electrodes had a slight decrease after 2 weeks' immersion in PBS and remained nearly unchanged in the next 6 weeks, indicating chemical stability in physiological environment.

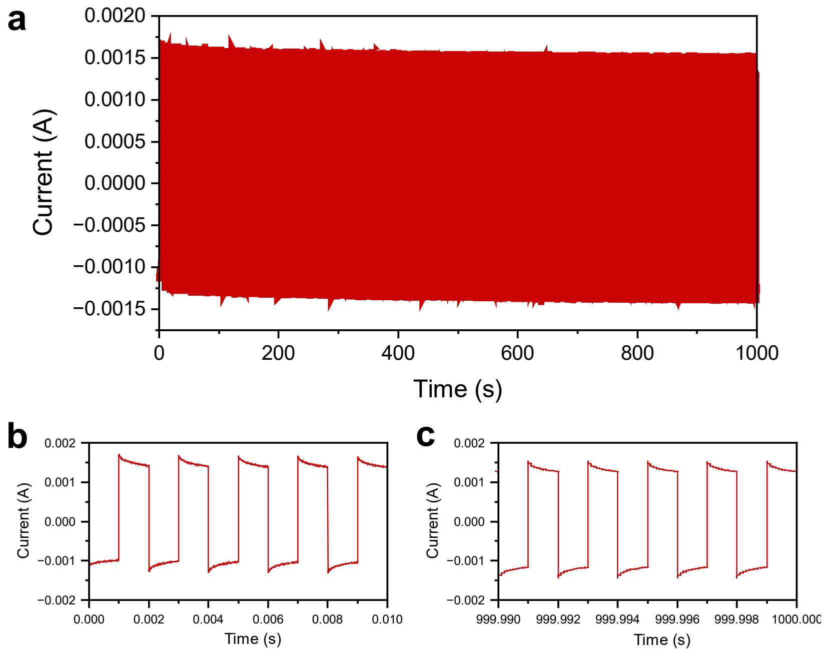


**Figure S10**. (a) Current-time curves of PEDOT fibers in bipolar pulse voltage stimulation measurement. The voltage amplitude is 0.5 V and the pulse duration for charge injection is 1 ms. (b-c) Current-time curves of PEDOT fibers in (b) the first 10 and (c) the last 10 pulse voltage stimulation cycles during one million of pulse voltage stimulation cycles. Again, it is found that the current-time curves of PEDOT fibers do not change after one million of pulse voltage stimulation cycles, indicating good electrochemical stability for use in neural stimulation.

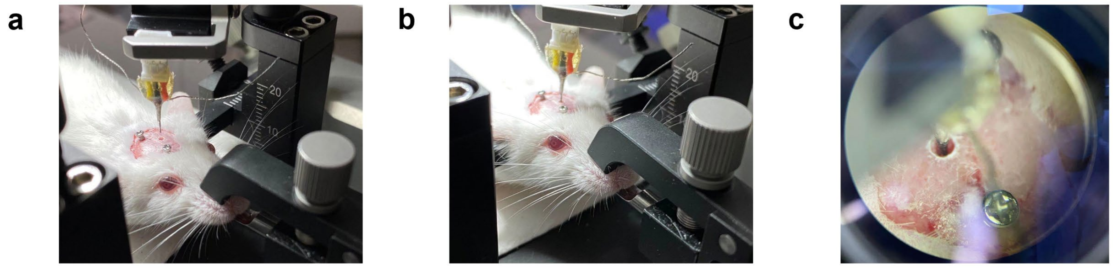


**Figure S11**. The implantation process of a maltose coated PEDOT microelectrode. (a) The PEDOT microelectrode was coated by maltose before implantation. (b) The coated maltose dissolved after microelectrode implantation within a few minutes. (c) The enlarged vision of the implanted PEDOT microelectrode.

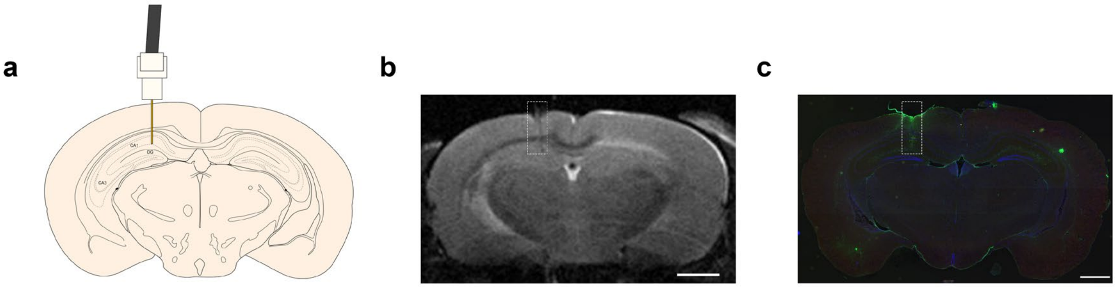


**Figure S12**. The implantation of PEDOT microelectrodes was confirmed by magnetic resonance imaging (MRI) and immunofluorescence staining. (a) A schematic diagram of the implantation site. (b) A MR image of the implanted microelectrodes. Scale bar: 1000 μm. (c) An immunofluorescence image after microelectrode implantation. Scale bar: 1000 μm.

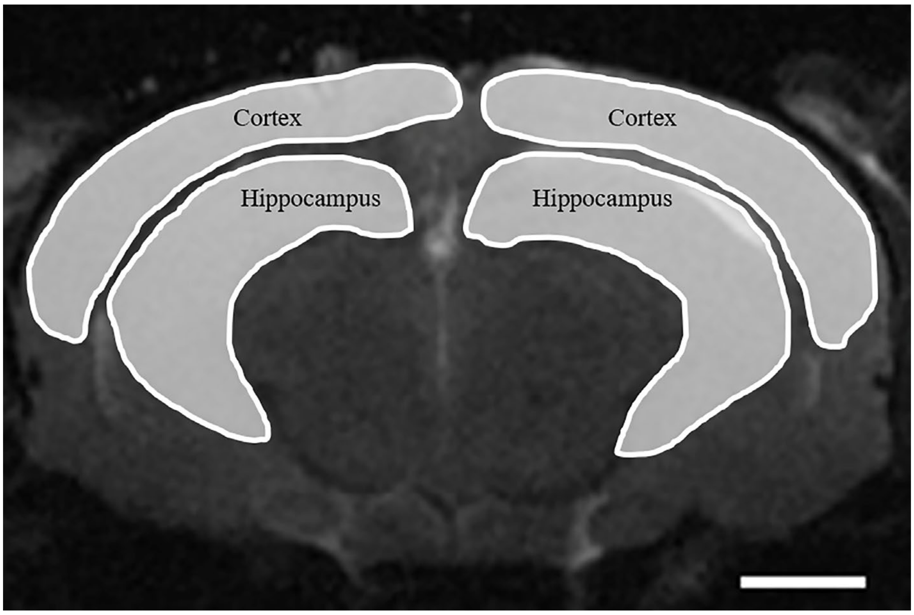


**Figure S13**. Regions of interest in mouse brain for the MRI analysis. Scale bar: 1000 μm.

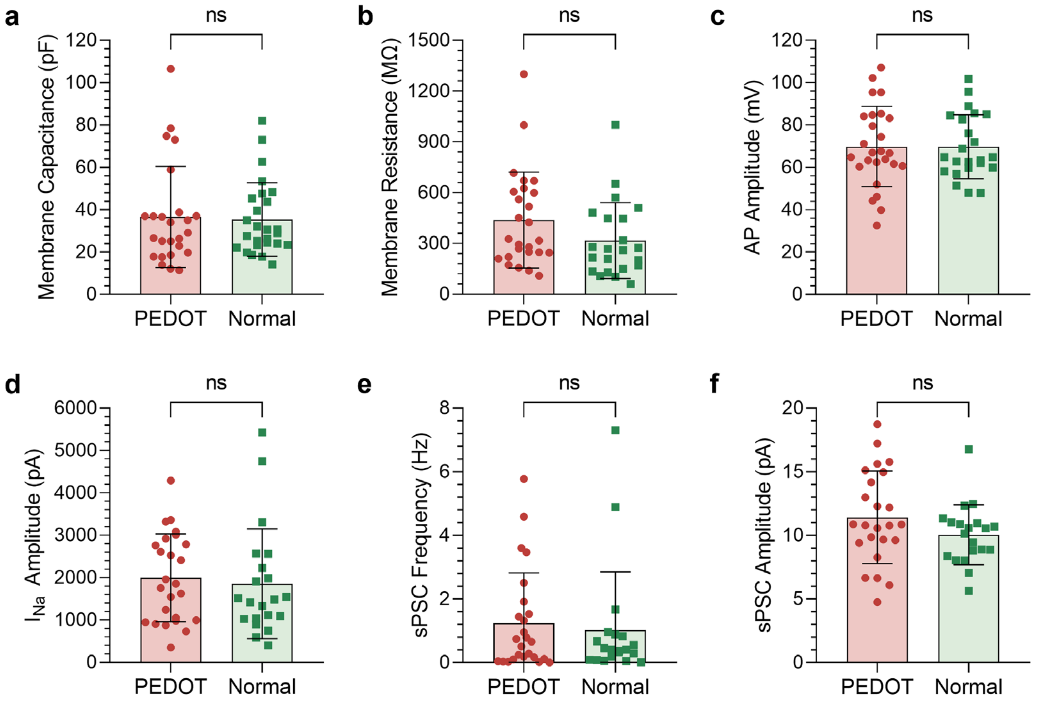


**Figure S14**. The comparison of electrophysiological properties between neurons of the bilateral hippocampi. (**a**, **b**) The measurements with (**a**) membrane capacitance ($n_1 = 25$ for the PEDOT group, $n_2 = 26$ for the normal group; student's independent-samples t tests, $t = 0.200$, $p > 0.05$) and (**b**) membrane resistance ($n_1 = 26$ for the PEDOT group, $n_2 = 22$ for the normal group; student's independent-samples t tests, $t = 1.603$, $p > 0.05$), two vital passive membrane electrical properties, showed no statistical differences between neurons of the bilateral sides. (**c**) By injecting depolarizing currents, neurons of both sides were able to generate action potential (AP), with no statistical differences of amplitudes ($n_1 = 26$ for the PEDOT group, $n_2 = 23$ for the normal group; student's independent-samples t tests, $t = 0.024$, $p > 0.05$). (**d**) Under the same clamping voltage, the amplitudes of the evoked voltage-gated sodium current ($I_{Na}$) exhibited no statistical differences between the bilateral hippocampi ($n_1 = 25$ for the PEDOT group, $n_2 = 21$ for the normal group; student's independent-samples t tests, $t = 0.417$, $p > 0.05$). (**e**, **f**) Spontaneous synaptic events were monitored comparably at the implantation and uninjured sides (Fig. 3i). The measurements with frequencies (n1 = 25 for the PEDOT group, n2 = 20 for the normal group; student's independent-samples t tests, t = 0.424, p > 0.05) and amplitudes ($n_1 = 24$ for the PEDOT group, $n_2 = 20$ for the normal group; student's independent-samples t tests, $t = 1.447$, $p > 0.05$) of spontaneous postsynaptic current (sPSC) were found no statistical differences between the bilateral hippocampi. * for $p < 0.05$, ** for $p < 0.01$, *** for $p < 0.001$ and *ns* for no statistic difference.

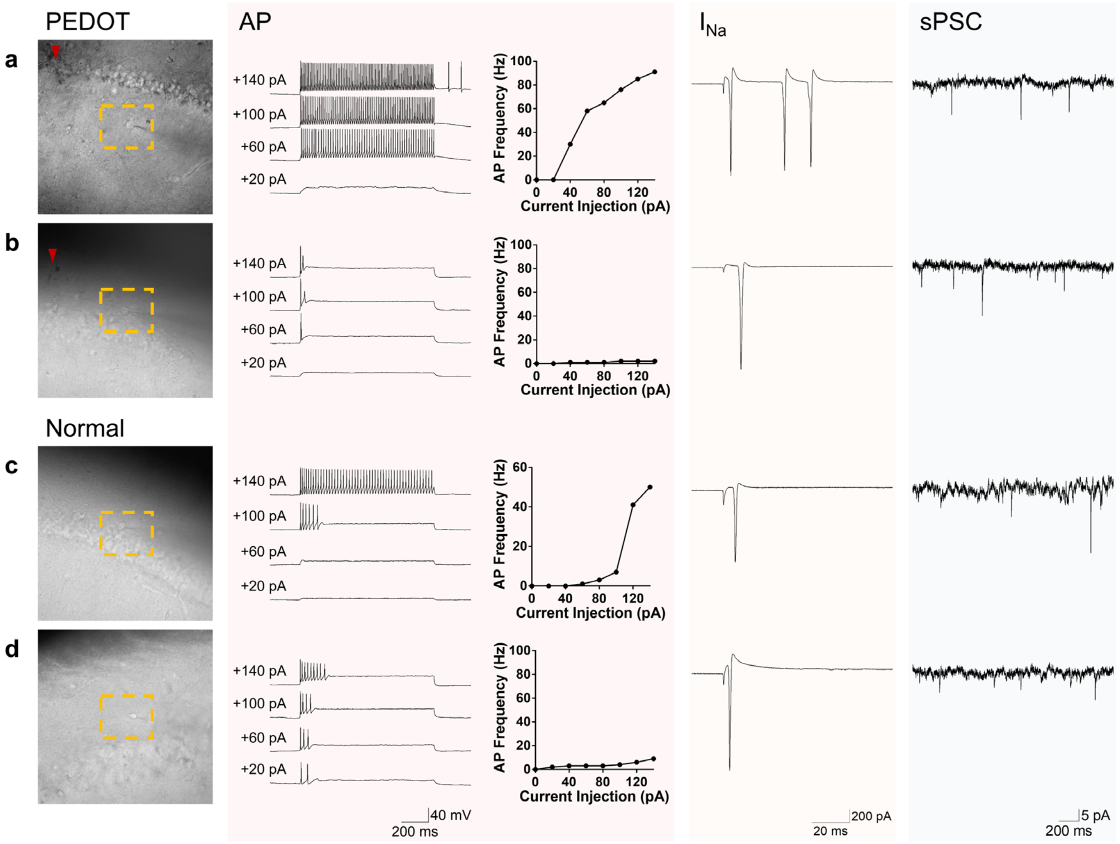


**Figure S15**. Specific examples of (a, c) excitatory neurons and (b, d) inhibitory neurons confirmed at (a, b) the implantation sides and (c, d) the uninjured sides. ▼ for the implantation trajectory.

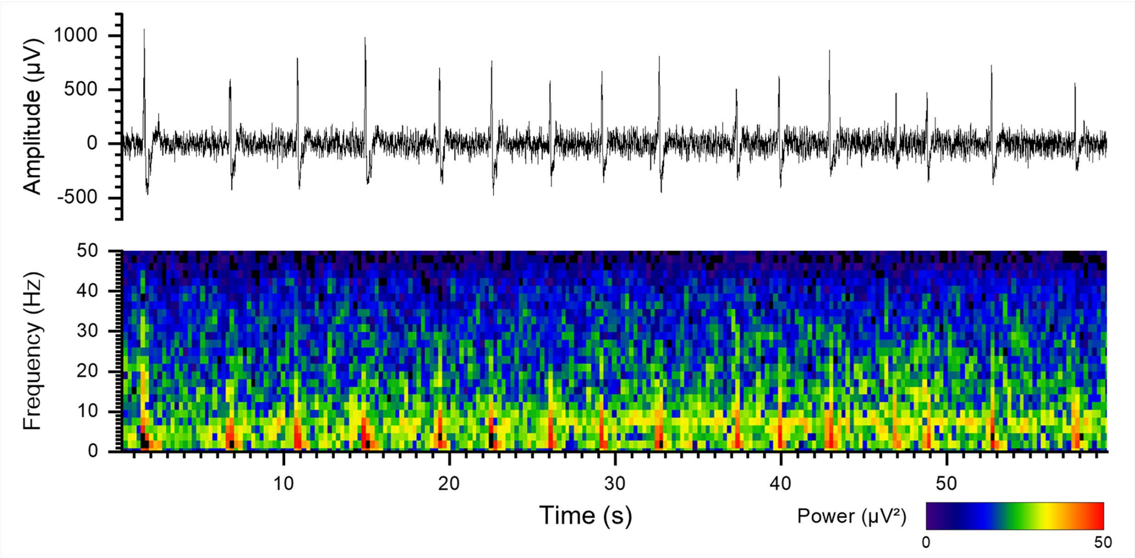


**Figure S16**. An example for epileptiform discharges recorded in the preictal state.

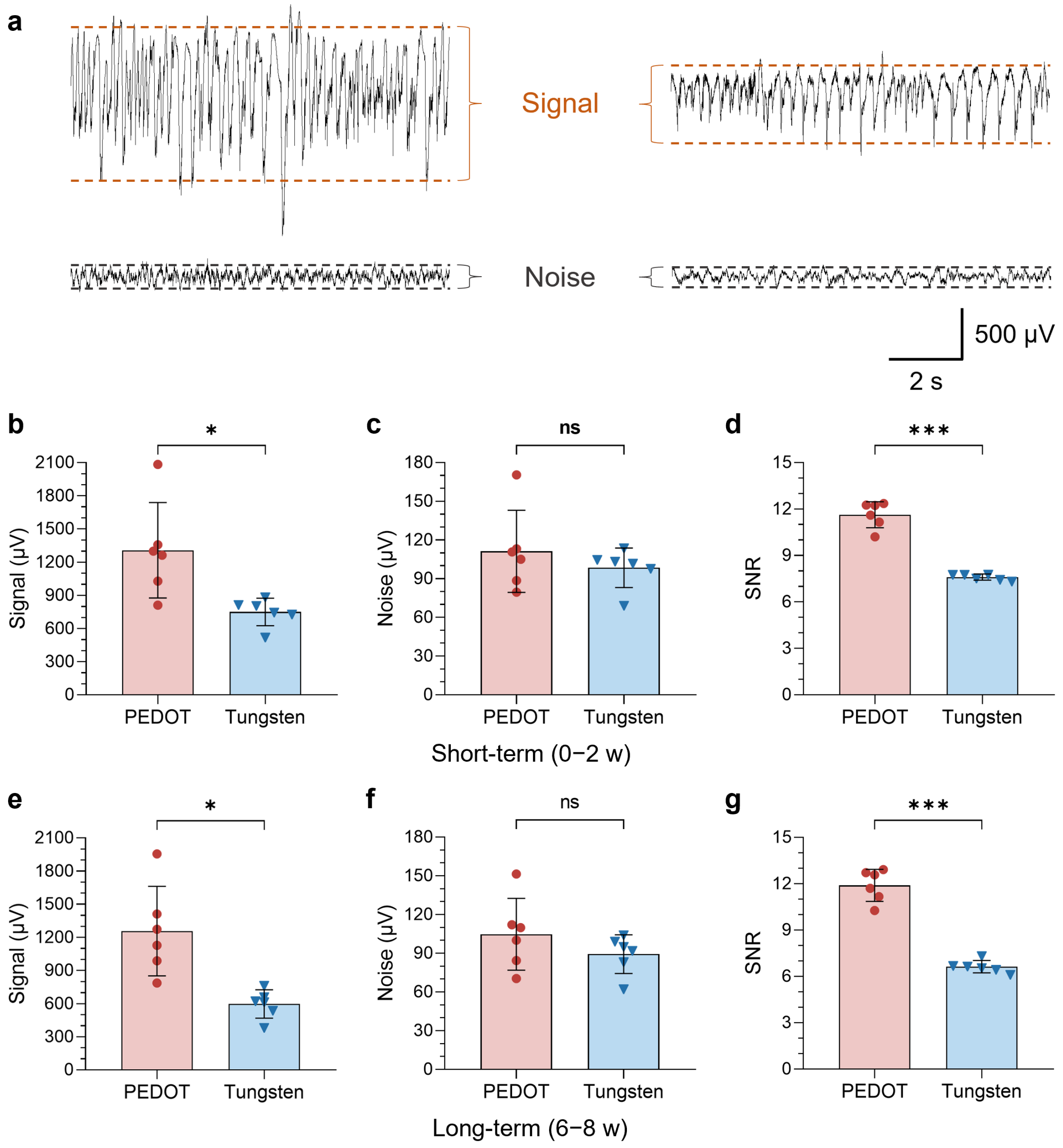


**Figure S17**. The signal and noise comparison between the PEDOT and the tungsten groups. (a) An illustration of how the signal amplitude and the noise level were calculated. (b-d) Quantitative comparisons of short-term (0−2 w) between two groups with (b) signal amplitude ($n = 6$ for each; student's independent-samples t tests, $t = 3.039$, $p < 0.05$), (c) noise level ($n = 6$ for each; student's independent-samples t tests, $t = 0.886$, $p > 0.05$), and (d) signal to noise ratio (SNR) ($n = 6$ for each; student's independent-samples t tests, $t = 11.455$, $p < 0.001$). (e-g) Quantitative comparisons of long-term (6−8 w) between two groups with (e) signal amplitude ($n = 6$ for each; student's independent-samples t tests, $t = 3.800$, $p < 0.01$), (f) noise level ($n = 6$ for each; student's independent-samples t tests, $t = 1.187$, $p > 0.05$), and (g) signal to noise ratio

(SNR) ($n = 6$ for each; student's independent-samples t tests, $t = 11.574$, $p < 0.001$). Compared with the tungsten, the signal amplitude in the PEDOT group was significantly higher, while the noise level didn't, thus leading to better SNR in both the short-term and long-term recordings.

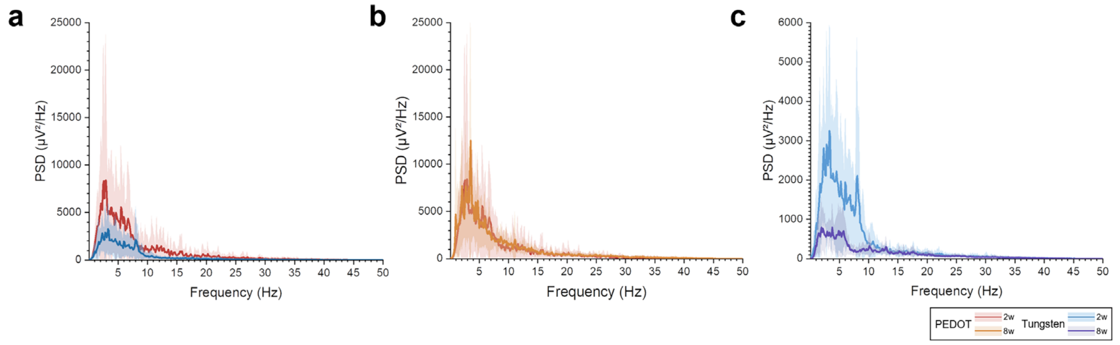


**Figure S18**. Power spectrum density (PSD) analyses for neural recordings (a) PSD comparison of short-term (0−2 w) recordings between the PEDOT and the tungsten groups. The PSD in the PEDOT group is significantly higher than that in the tungsten group at the frequency range between 0−30 Hz. (b, c) PSD comparison between short-term (0−2 w) and long-term (6−8 w) recordings in (b) the PEDOT and (c) the tungsten groups respectively. The PSD in the PEDOT group present similar levels in both the short-term and long-term recordings, while the PSD in the tungsten group significantly damps in the long-term recordings.

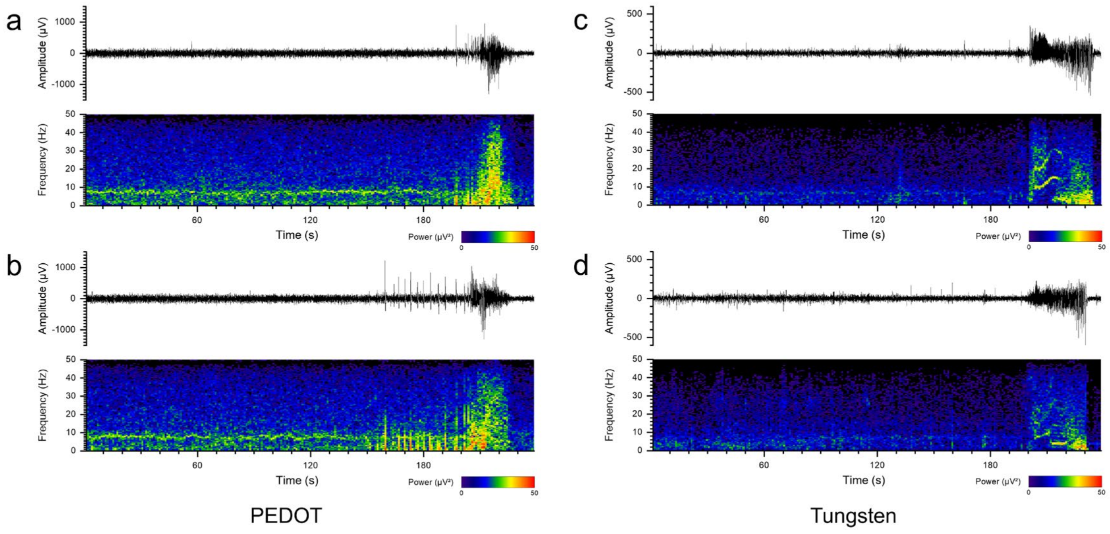


**Figure S19**. Examples for (a, c) the short-term (0−2 w) and (b, d) the long-term (6−8 w) recordings of epileptic seizures acquired by different microelectrodes.

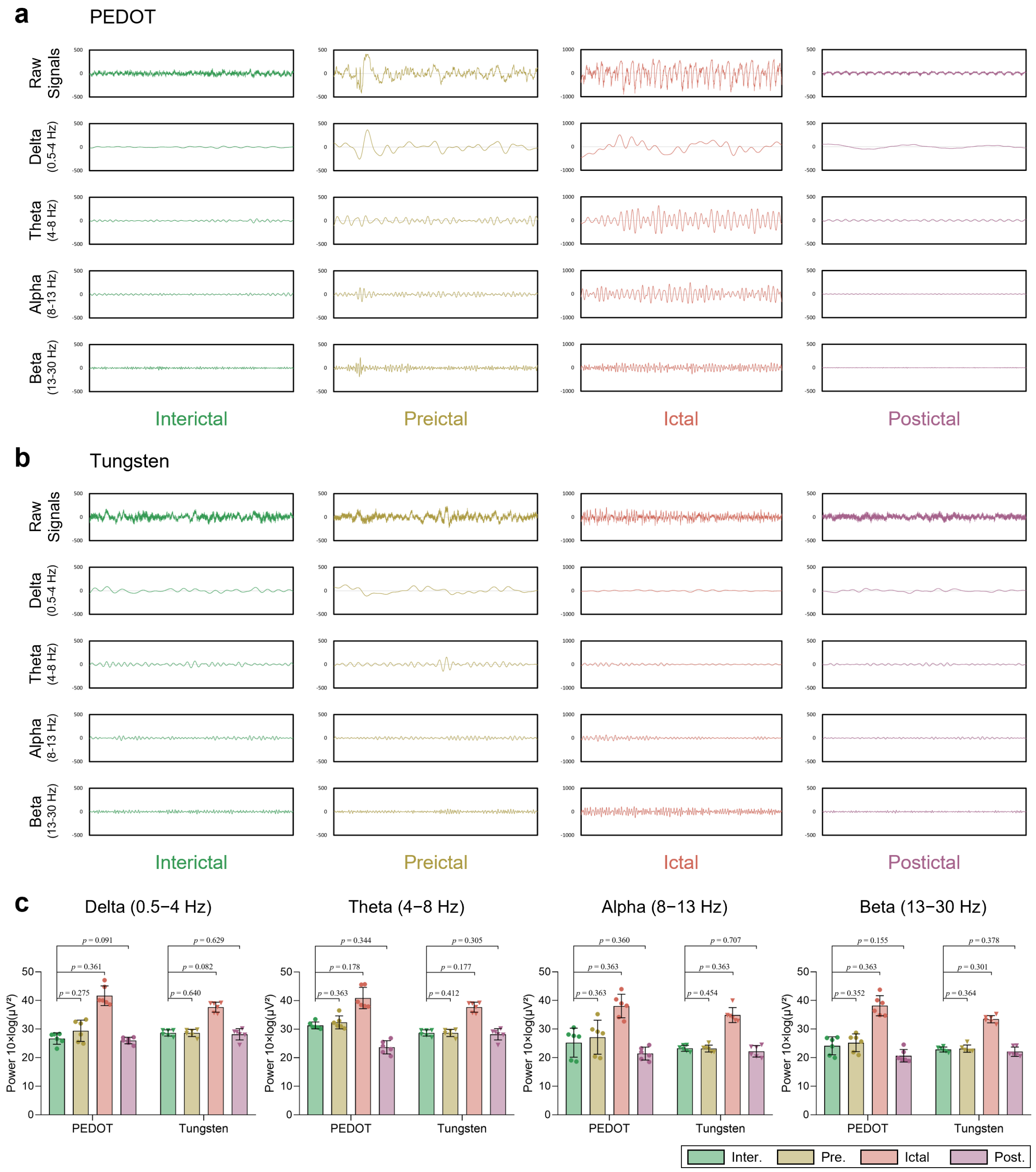


**Figure S20**. The signal details of four epileptic states. (a, b) The digital filtering signals across different frequency bands with delta range (0.5−4 Hz), theta range (4−8 Hz), alpha range (8−13 Hz), and beta range (13−30 Hz). (a) for the PEDOT group and (b) for the tungsten group; signal amplitude: μV; time window: 5 s. (c) The band power of four epileptic states across different frequency bands ($n = 6$ for each; student's paired-samples t tests between the interictal and the other states in each frequency band). It can be seen that the signal waves of four epileptic states exhibit more obvious distinctions in the PEDOT group, compared with the tungsten group.

Similarly, the differences of band power between states are also more significant in the PEDOT group.

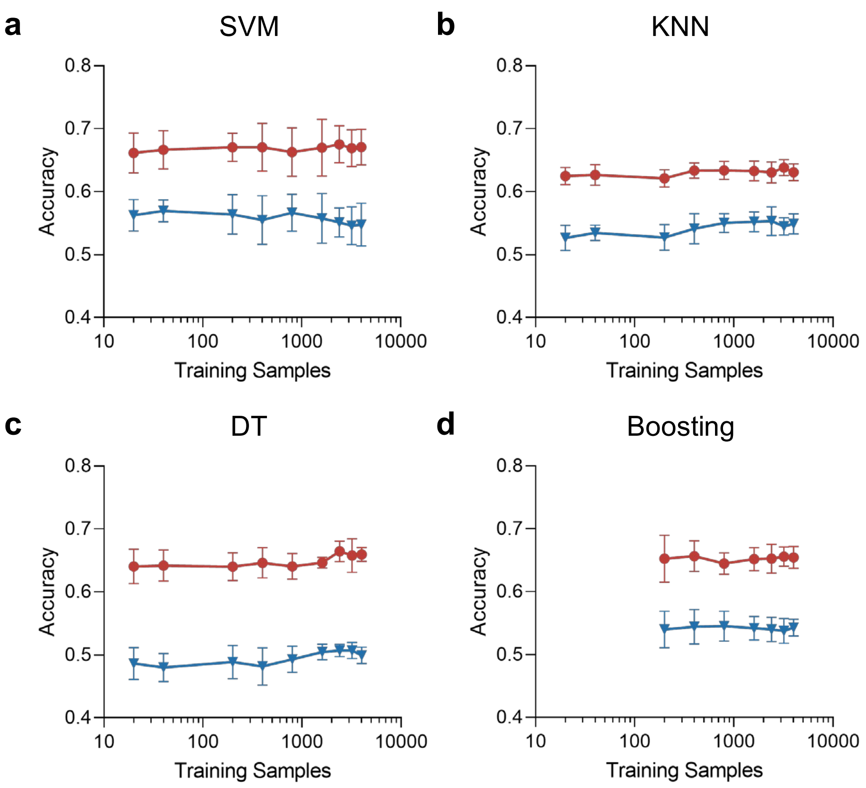


**Figure S21**. (a-d) Classification accuracy as a function of training samples (mean ± STD) for the conventional machine learning algorithms, including (a) support vector machine (SVM), (b) K nearest neighbor (KNN), (c) decision tree (DT), and (d) boosting respectively. It can be seen that the performance of four algorithms exhibits no dependence on the amount of training data. Notably, boosting was failed to be trained when the amount of training samples was below 100.

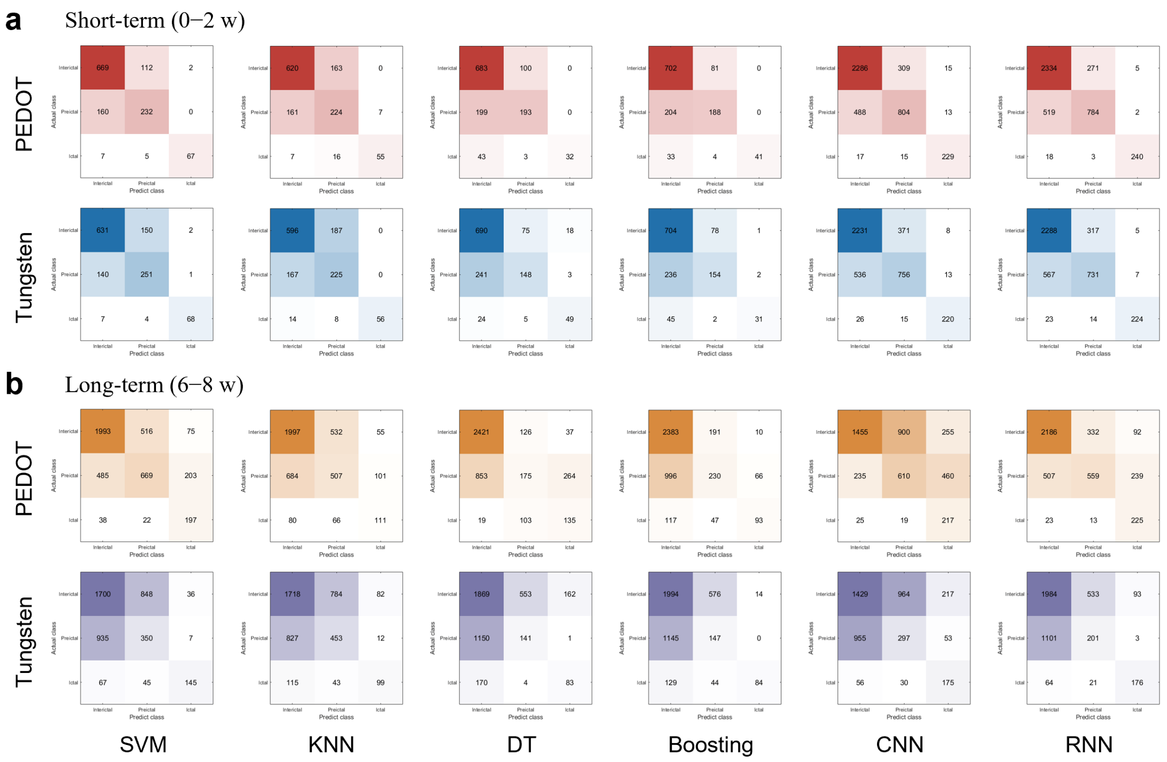


**Figure S22**. Confusion matrixes for epileptic state classification by using different algorithms in (a) the short-term (0−2 w) and (b) the long-term (6−8 w) recordings respectively. Compared with the tungsten group, the amounts of correct predictions in the PEDOT group were larger in both the short-term and long-term recordings, especially in the interictal and the preictal classes. The incorrect predictions in the long-term recordings mainly increased in the preictal classes, which might be attributed to the harder distinguish of the preictal signals from the interictal signals after SNR decreasing. This situation was worse in the tungsten group, and caused more significant degradation in its long-term recordings.

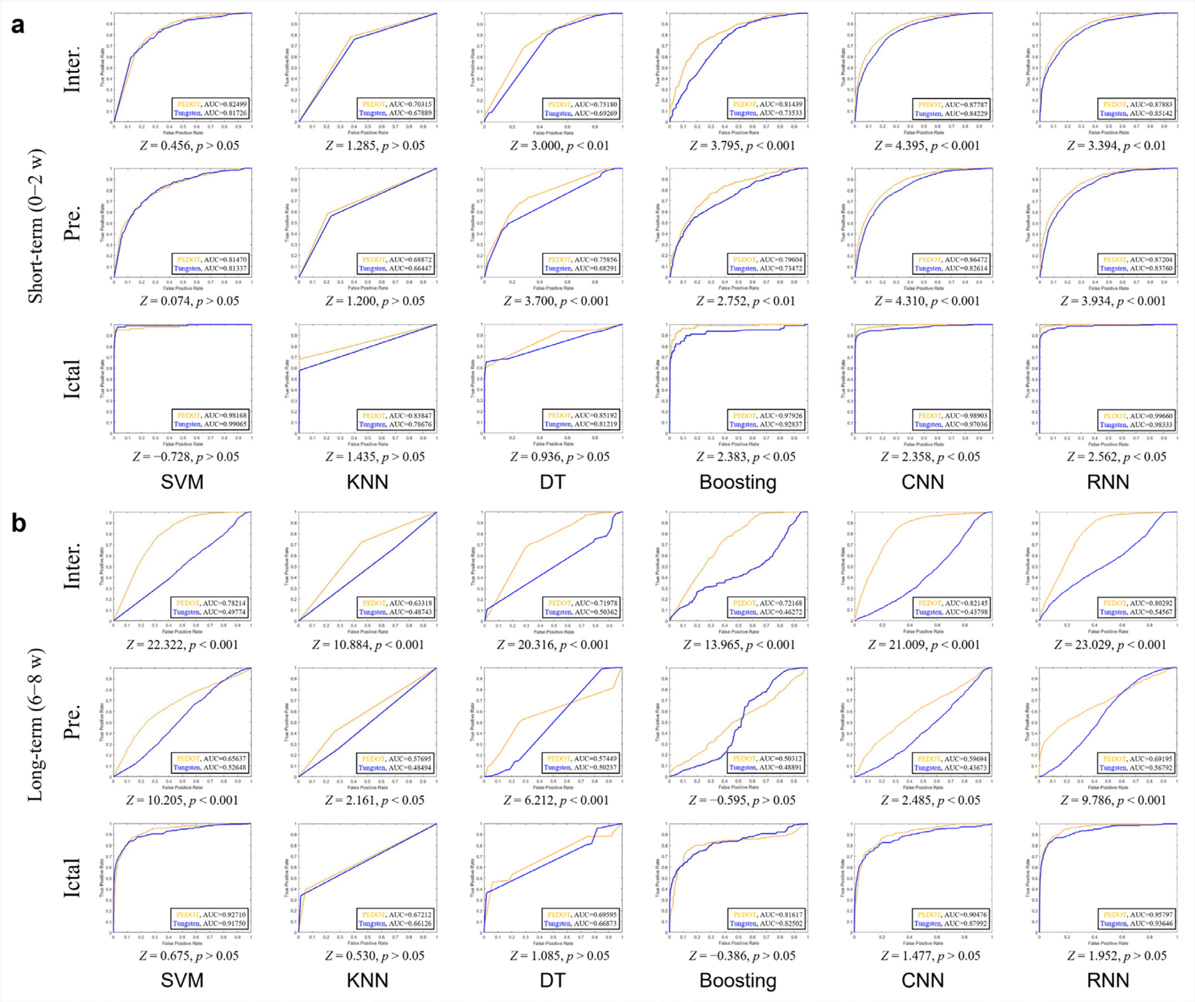


**Figure S23**. Receiver operating characteristic (ROC) curve for epileptic state classification by using different algorithms in (a) the short-term (0–2 w) and (b) the long-term (6–8 w) recordings respectively. For most algorithms, the areas under the ROC curve (AUC) of the interictal and preictal classes are significantly higher in the PEDOT group than those in the tungsten group, while the AUC of the ictal class show no significant differences between groups. Compared with the short-term recordings, the differences of AUC between two groups were more significant in both the interictal and the preictal classes. The evidence suggests that the better SNR of the PEDOT group is conducive to distinguish the preictal signals from the interictal signals.

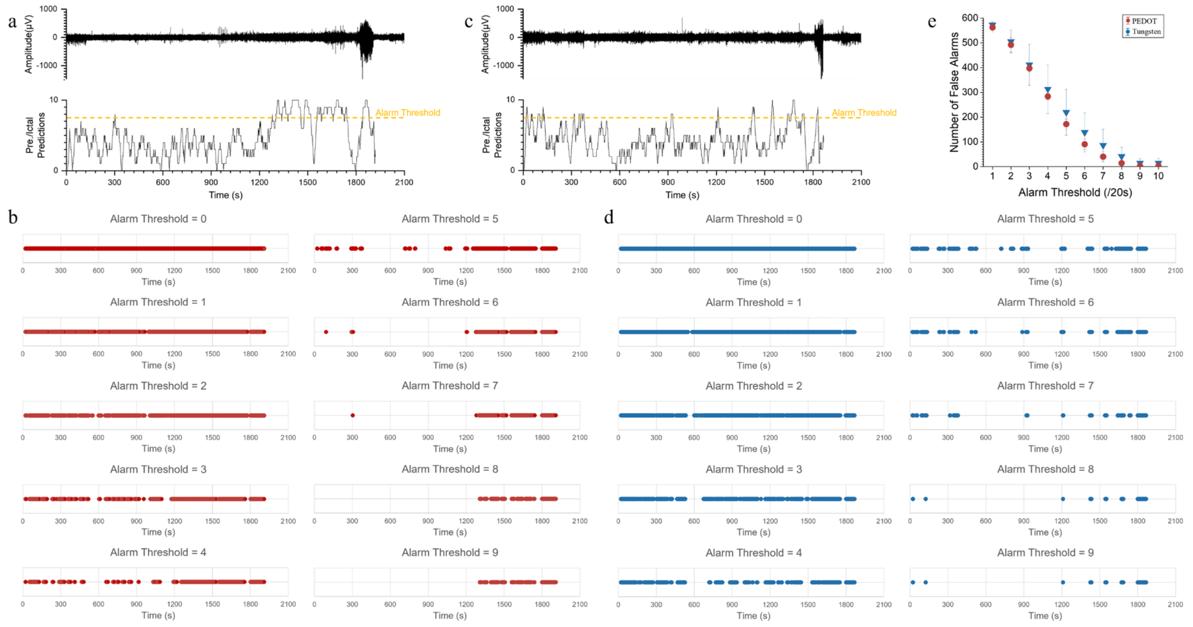


**Figure S24**. (a-d) A gradient of alarm threshold and its practical effects in the PEDOT (a, b) and the tungsten (c, d) groups respectively. Alarms were triggered when the number of preictal or ictal predictions in the past 20 seconds crossed the decision threshold. The ictal state (seizure) begins at time 1800 s, and the periods of 600 s (10 min) before seizure onset are defined as the preictal state. (e) The number of false alarms along with the threshold gradient. It can be found that the number of false alarms was very few when the threshold was over 7, and the PEDOT group presented the less false alarms in the interictal periods and the earlier warnings in the preictal periods.

**Table S1. The statistical results for classification performance in short-term recordings**

| Classifier | Group | ACC | SPE | | | SPN | | |
|---|---|---|---|---|---|---|---|---|
| | | | Inter. | Pre. | Ictal | Inter. | Pre. | Ictal |
| SVM | PEDOT | 0.795 | 0.867 | 0.640 | 0.846 | 0.683 | 0.878 | 0.997 |
| | Tungsten | 0.752 | 0.811 | 0.625 | 0.795 | 0.666 | 0.828 | 0.995 |
| KNN | PEDOT | 0.718 | 0.792 | 0.571 | 0.705 | 0.643 | 0.792 | 0.994 |
| | Tungsten | 0.700 | 0.761 | 0.574 | 0.718 | 0.615 | 0.774 | 1.000 |
| DT | PEDOT | 0.725 | 0.872 | 0.492 | 0.410 | 0.485 | 0.880 | 1.000 |
| | Tungsten | 0.708 | 0.881 | 0.378 | 0.628 | 0.436 | 0.907 | 0.982 |
| Boosting | PEDOT | 0.743 | 0.897 | 0.480 | 0.526 | 0.496 | 0.901 | 1.000 |
| | Tungsten | 0.710 | 0.899 | 0.393 | 0.397 | 0.402 | 0.907 | 0.997 |
| CNN | PEDOT | 0.795 | 0.876 | 0.616 | 0.877 | 0.678 | 0.887 | 0.993 |
| | Tungsten | 0.768 | 0.855 | 0.579 | 0.843 | 0.641 | 0.866 | 0.995 |
| RNN | PEDOT | 0.804 | 0.894 | 0.601 | 0.920 | 0.657 | 0.905 | 0.998 |
| | Tungsten | 0.777 | 0.877 | 0.560 | 0.858 | 0.623 | 0.885 | 0.997 |

**Table S2. The statistical results for classification performance in long-term recordings**

| Classifier | Group | ACC | SPE | | | SPN | | |
|---|---|---|---|---|---|---|---|---|
| | | | Inter. | Pre. | Ictal | Inter. | Pre. | Ictal |
| SVM | PEDOT | 0.714 | 0.772 | 0.604 | 0.673 | 0.687 | 0.791 | 0.973 |
| | Tungsten | 0.548 | 0.723 | 0.207 | 0.506 | 0.294 | 0.744 | 0.988 |
| KNN | PEDOT | 0.633 | 0.773 | 0.392 | 0.432 | 0.507 | 0.790 | 0.960 |
| | Tungsten | 0.549 | 0.665 | 0.351 | 0.385 | 0.392 | 0.709 | 0.976 |
| DT | PEDOT | 0.661 | 0.937 | 0.135 | 0.525 | 0.383 | 0.949 | 0.922 |
| | Tungsten | 0.506 | 0.723 | 0.109 | 0.323 | 0.148 | 0.804 | 0.958 |
| Boosting | PEDOT | 0.655 | 0.922 | 0.178 | 0.362 | 0.281 | 0.916 | 0.980 |
| | Tungsten | 0.538 | 0.772 | 0.114 | 0.327 | 0.178 | 0.782 | 0.996 |
| CNN | PEDOT | 0.547 | 0.557 | 0.467 | 0.831 | 0.834 | 0.680 | 0.817 |
| | Tungsten | 0.455 | 0.548 | 0.228 | 0.670 | 0.354 | 0.654 | 0.931 |
| RNN | PEDOT | 0.711 | 0.838 | 0.428 | 0.862 | 0.662 | 0.880 | 0.915 |
| | Tungsten | 0.565 | 0.760 | 0.154 | 0.674 | 0.256 | 0.807 | 0.975 |
